# Subjective understanding of actions and emotions requires interaction of the semantic and action observation networks

**DOI:** 10.1101/2021.04.15.439961

**Authors:** Minye Zhan, Rainer Goebel, Beatrice de Gelder

## Abstract

Previous studies have investigated the neural basis of action and emotion perception conveyed by whole body postures and movements. However, subjective understanding of other people’s bodily actions and emotions, raises different issues that are currently not well understood and have so far seldom been investigated. In this 7T fMRI study, we proceeded beyond conventional univariate/multivariate analyses with predefined categories, and examined the representational geometry of subjective action- and emotion-understanding by mapping individual subjective reports with word embeddings. Dimensionality reduction revealed that the representations for perceived action and emotion were high dimensional, each correlated to but were not reducible to the predefined action and emotion categories. Searchlight representational similarity analysis showed that the representations in the left middle superior temporal sulcus and left dorsal premotor cortex corresponded to the subjective action and emotion understanding of the participants. Furthermore, using task-residual functional connectivity and hierarchical clustering, we found that areas in the action observation network and the semantic/default-mode network were functionally connected to these two seed regions, and showed similar representations. Our study provides direct evidence that both networks were concurrently involved in subjective action and emotion understanding. Our approach shows how subjective understanding of action and emotion stimuli can be reliably studied and complement insights based on from predefined stimulus categories.

## Introduction

Imagine you are walking into a large conference hall and you see a person waving in your direction from far away. Within hundreds of milliseconds, this triggers a cascade of implicit and explicit brain processes: “Do I know this person? Is she waving towards me, perhaps to notify me something? Since I don’t know her and she seems to be quite happy, is she perhaps waving towards someone close by who might be her acquaintance?” This is an example of the multifaceted information about the action, emotion, intention and identity that human bodies routinely convey, which we process effortlessly in daily life. Our brains extract and combine those various dimensions of information, enabling us to reach a personal/subjective understanding, and to act upon it.

These different dimensions of information have been the focus of specialized lines of inquiry. Univariate studies of action observation have found a fronto-parietal network (action observation network) including the intraparietal sulcus (IPS) and the dorsal and ventral premotor areas (PMd, PMv)(Caspers et al., 2010; Grafton and Hamilton, 2007; Rizzolatti et al., 2014). Studies focused on the body form found body/body parts-sensitive regions in the ventral-lateral pathway, the extrastriate body area (EBA) and the fusiform body area (FBA) (Peelen and Downing, 2007), while the posterior superior temporal sulcus (pSTS) is sensitive to the biological motion of both faces and bodies (Allison et al., 2000). Studies investigating how body posture and movements convey emotional information reported activation in areas associated with visual form, action and movement perception, which goes hand in hand with activation in more emotion-related areas including the IFG, insula and subcortical structures (de Gelder, 2006; de Gelder et al., 2004; Dricu and Frühholz, 2016; Kober et al., 2008; Lindquist et al., 2012; Molenberghs et al., 2012; Sinke et al., 2010).

So far, there has been limited evidence that bodily action and emotion processing also involves the default mode network (DMN) areas (Andrews-Hanna, 2012; Buckner et al., 2008), including the temporo-parietal junction (TPJ), precuneus, dorsal/ventral medial prefrontal cortex (dmPFC, vmPFC) (Amodio and Frith, 2006; Saxe et al., 2006). Univariate activation of DMN areas was rarely found in action perception, apart from a few studies (e.g. Brass et al., 2007; de Lange et al., 2008). Therefore the DMN was generally considered as a separate system that does not interact with the action observation network (Van Overwalle and Baetens, 2009). However, instead of univariate activation, studies using multivariate methods such as representational similarity analysis (RSA) (Kriegeskorte et al., 2008; Nili et al., 2014) found that the DMN areas were involved in more general and abstract emotion and valence processing, for some stimuli types other than bodies (Chikazoe et al., 2014; Peelen et al., 2010; Skerry and Saxe, 2015).

Most previous fMRI studies presented participants with exemplars from a few predefined categories, and often used an explicit action or emotion recognition/categorization task, serving as a proxy for participants’ subjective action and emotion understanding. This approach reflects the traditional view of a few basic emotions (Ekman, 1999), assuming that (1) there exist a few discrete categories of actions and emotions that participants are routinely able to recognize; (2) there is high inter-individual consistency in recognizing and identifying these categories.

Under these assumptions, individual variability is typically treated as noise, partially due to the difficulty to objectively quantify it across participants, especially for the case of verbal reports. When individual variability is ignored, neural substrates that do not conform to the predefined responses or the averaged behavior were missed. However, individual variability is prevalent in multiple functions from perception to cognition, and some individual variability can show stable brain-behavior mappings, including the size or function of specific brain areas (Charest et al., 2014; Kanai and Rees, 2011; Seghier and Price, 2018). Specifically concerning action and emotion perception, individual variability is seen in multiple validation studies for facial expressions (e.g. Goeleven et al., 2008; Langner et al., 2010). Other studies also suggested that different observers may reach a different understanding of the action/emotion, depending on factors such as the personality of the viewer (Van den Stock et al., 2015) and various contexts (Aviezer et al., 2008; Kret and de Gelder, 2010; Righart and de Gelder, 2006).

Recently, researchers argued that studying subjective experiences is important for understanding high-level cognitions such as emotion, language, and music (Hartley and Poeppel, 2020; LeDoux and Hofmann, 2018). The presence of subjective variability in emotional perception and production is also acknowledged in the recent literature (Barrett et al., 2019; Cowen and Keltner, 2021). In response to these challenges, new studies utilized very large stimulus sets, subjective reports, and the associated semantic space (Cowen et al., 2019; Cowen and Keltner, 2021, 2020, 2017). The results found a large amount (13 to 28) of emotional categories could be observed in subjective reports for facial, vocal, musical emotional expressions. There were no discrete category boundaries, and the semantic space of these categories was also high-dimensional. The brain-behavior mappings of such rich emotion representations seen in subjective experience independent from the predefined basic categories are still largely uncharted.

With the advancement of deep neural networks and natural language processing techniques, word embeddings (see Boleda, 2020 for a review) are now increasingly used to describe semantic concepts in an objective and quantitative way, and recent studies started linking them to brain activity (Hebart et al., 2019; Zhang et al., 2020).

In this high-resolution 7T fMRI study (1.2 mm isotropic resolution, voxel volume = 1.728 mm^3^) with RSA (Kriegeskorte et al., 2008; Nili et al., 2014), we aimed to go beyond conventional analysis with predefined categories, and to examine the shared neural systems that enabled each participant to independently reach his/her own version of action/emotion understanding, be it a different version or not from other participants. To ensure subjective perception, the participants were not given any action/emotion category labels or categorization task. During functional runs, 10 participants passively viewed a large stimulus set of 10 predefined bodily action categories (6 neutral, 4 emotional categories; each participant viewed 40 stimuli). Immediately following the scanning session, participants provided free reports of the subjectively perceived action and emotion for each stimulus.

We first examined the representational geometry of the subjective reports by principal component analysis (PCA) and RSA, after mapping all subjective report entries into a common 300-dimensional vector space using Deconf word embeddings (Pilehvar and Collier, 2016). In the conventional analyses of predefined categories, we examined their neural representations with RSA searchlight and RSA regression. We also estimated the body joint positions using the OpenPose library (Cao et al., 2019), to account for areas related to processing of low- and mid-level visual features. In analyses examining representations of subjective understandings, we again searched for corresponding neural representations by RSA searchlight, but with individualized model matrices. For the resulting group-level areas, we then examined their putative direct upstream/downstream areas by task-residual functional connectivity and hierarchical clustering. We found neural representations for both perceived action and perceived emotion, and the two analyses with predefined categories and with the subjective reports converged as they both indicated joint involvement of the action observation network and the DMN/semantic network, noting that the analyses of the subjective understanding provided much more direct evidence.

## Results

### RDM construction

In the scanning session, participants passively viewed 40 gray-scale images from one of two balanced stimuli sets (**Fig. 1A**. See **Fig. S1** for the complete sets of stimuli), to have enough repetitions per stimulus while having enough stimulus image variability for each participant. We used a slow event-related design, where each image (2.60 × 4.26 degrees) was presented for 500 ms, followed by an inter-stimulus interval of either 7.5, 9.5 or 11.5 s. Each image was presented 12 times.

**Figure 1.**
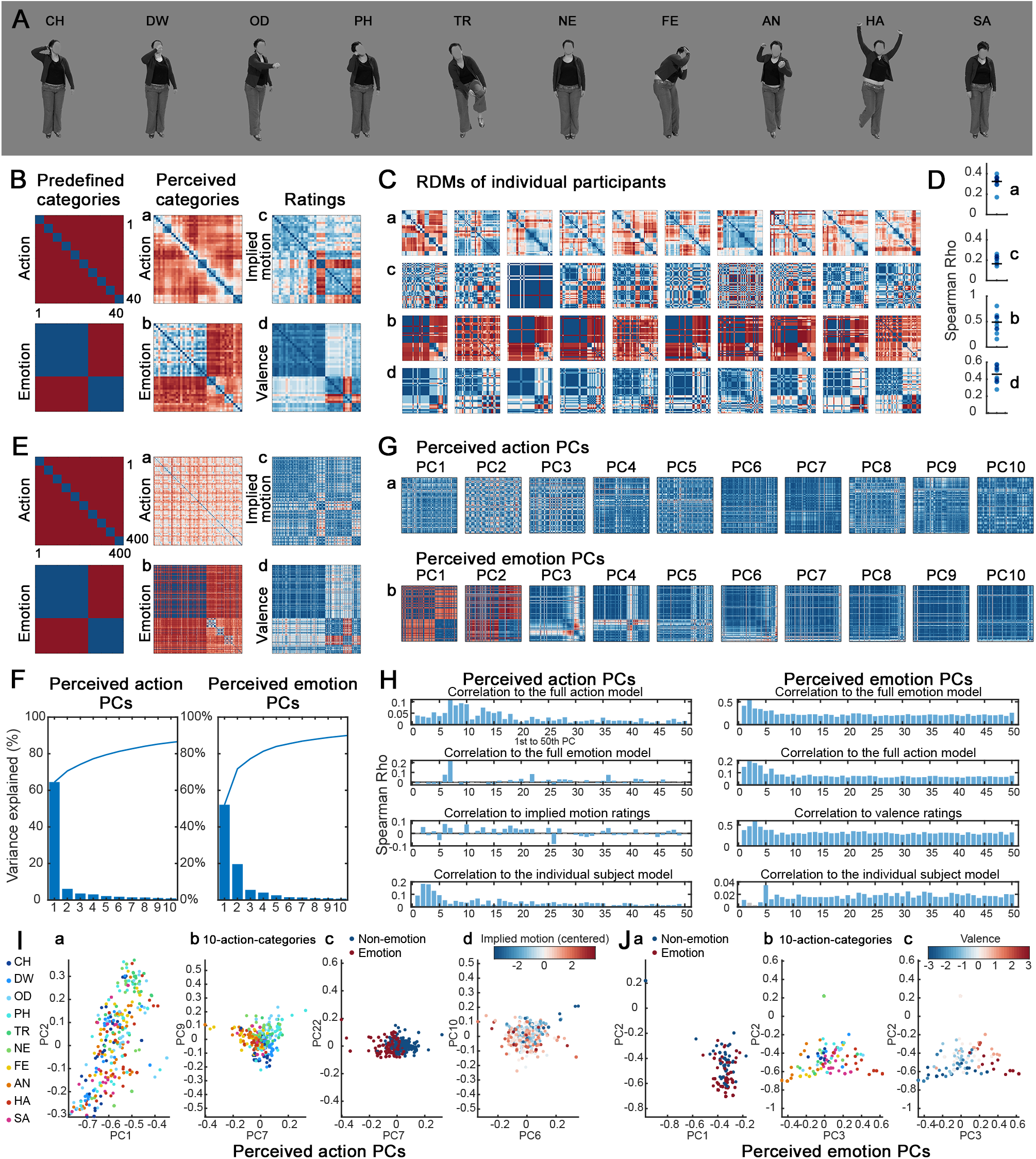
The representational geometries of subjective reports were high-dimensional, and showed considerable individual variability. **A**. Stimuli examples of 10 action categories, performed by one of 8 actors. Category abbreviations: CH: combing hair; DW: drinking water; OD: opening door; PH: phone; TR: putting on trousers; NE: neutral standing still; FE: fearful; AN: angry; HA: happy; SA: sad. **B**. RDMs for predefined categories, perceived categories, ratings, averaged across 10 participants (40 by 40 matrices, sorted by the 10 action categories). **a**. perceived action; **b**. perceived emotion; **c**. rated implied motion; **d**. rated valence. Colors in **B, C, E, G** were scaled automatically by the minimal/maximal values within each RDM. Blue: similar; color: dissimilar. **C**. RDMs for individual participants (each column) showed considerable individual variability. RDM types **a** to **d** correspond to **a** to **d** in **B**. The left 5 participants viewed stimulus set A, the other 5 viewed set B. **D**. Correlation to the predefined models for individual participants were not high. **a** to **d** correspond to the ones in **B** and **C**. Black bars: group average. **E** to **J**. PCA results of all 400 behavioral report items for perceived action and perceived emotion. The smaller PCs of the perceived action and the first PCs of the perceived emotion showed correspondence to the predefined RDMs. **E**. Same to **B**, but for all behavioral report items across participants (400 by 400 matrices). **F**. The first 10 principal components (PCs) explained a large amount of variance. Curves: cumulative explained variance (%). **G**. RDMs for scores of the first 10 PCs (Euclidean distance), for perceived action (**a**) and perceived emotion (**b**). **H**. RDMs for the first 50 PCs, correlated to the predefined action, emotion models, behavioral rating models, and to the individual subject model. The first PCs of the perceived action correlated more to the individual subject model. Blue bars: FDR q<0.05. **I** and **J**. Perceived action (**I**) and emotion (**J**) report items, plotted against the first two PCs, and against the two PCs with highest correlation coefficients to each of the models.

In the behavioral session directly after the fMRI scan, participants reported their subjective understanding of the action and emotion of each stimulus by typing a short description, and rated the implied motion and valence on a scale of 1 to 7 (See **Table S1** for the exact questions, and examples of participants’ free reports).

Individual reports were analyzed by mapping all response entries to the same high-dimensional space using the pre-trained Deconf word embeddings (Pilehvar and Collier, 2016), which combined the word2vec word embeddings and the WordNet database (see Methods for details). Specifically, we lemmatized all the verbs, nouns, adjectives, adverbs typed in by the participants (e.g. fighting/fights→fight), and selected the corresponding meaning for each word in WordNet 3.1 (https://wordnet.princeton.edu/). We then retrieved the corresponding 300-dimensional vectors from Deconf embeddings, averaged the word vectors in each response entry, and computed RDMs for perceived action and perceived emotion (cosine distance) for each individual participant, see **Fig. 1Bab** and **1Cab**.

Implied motion and valence ratings were one-dimensional attributes provided by individual participants, which were related to but not fully represent the action and emotion aspects of each stimulus. We used them to complement the analysis of action- and emotion-understanding. These RDMs were computed directly from the individual behavioral ratings (Euclidean distance, **Fig. 1Bcd, Ccd**).

For predefined categories (10 action categories, non-emotional/emotional actions), we followed the RSA and perceptual categorization literature and constructed model RDMs (Euclidean distance, **Fig. 1B**, first column), assuming that the stimuli representations were similar within categories, but were different across categories between stimuli (Freedman et al., 2001).

We examined the correlations of the subjective report and the rating RDMs with the two predefined RDMs. Throughout the whole study, correlations between RDMs were computed with Spearman correlation, and submitted to a one-sample t test against 0 (two-tailed) at the group level, after Fisher’s Z transformation.

### Perceived categories largely corresponded to the predefined ones, but with considerable individual variability

All four types of subjective report and rating RDMs were significantly correlated to the predefined ones (one-sample t test against 0; coefficient of variation, CV, showing inter-individual variability), perceived action to predefined action RDMs: mean rho=0.324, p=8.55×10^−8^, CV=20.4%; perceived emotion to predefined emotion RDMs: mean rho=0.491, p=0.000164, CV=51.2%; rated implied motion to the predefined action RDMs: mean rho=0.166, p=0.000541, CV=60.4%; rated valence to the predefined emotion RDM: mean rho=0.496, p=5.91×10^−7^, CV=25.5%. This indicates that the individual subjective understanding largely corresponded to the predefined categories, especially for emotions, although there was considerable individual variability (**Fig. 1C, D**). For all four types of subjective reports and ratings, the two stimuli sets did not result in set-specific reports: the Spearman correlation similarity across individual RDMs within a stimuli set was not different from the similarity between sets (Wilcoxon rank sum test, all p>0.53).

We then examined the individual variability, to assess how appropriate it is to use group-averaged RDMs to perform further analyses in individual participants. We first obtained a group-averaged RDM from all 10 individual RDMs, then computed its Spearman correlation to each individual RDM (one-sample t test against 0; CV). When inter-individual consistency is high, the group-averaged RDM should have high correlation to each individual RDM, and a low CV. For valence, the group-averaged RDM was very consistent with individual RDMs: mean rho=0.900, p=6.91×10^−8^, CV=19.9%. However, the other 3 group-averaged behavioral RDMs were less consistent with individual RDMs, again showing considerable individual variability: perceived action RDM: mean rho=0.517, p=4.74×10^−6^, CV=32.7%; perceived emotion RDM: mean rho=0.565, p=6.02×10^−5^, CV=44.9%; rated implied motion RDM: mean rho=0.621, p=8.71×10^−5^, CV=47.1%. This moderate consistency is unlikely to be caused by our small sample size, but seems to be a persisting effect regardless of the sample size. In validation studies of facial emotion stimuli with much larger sample sizes (Goeleven et al., 2008; Langner et al., 2010), the consistency (measured by reporting accuracies) was in a similar range as the current study (see **Table S2** for comparison).

With these moderate correlation coefficients, indicating less tight brain-behavior links for individual participants, using group-averaged RDMs or the predefined category RDMs in subsequent analyses may be less suitable to capture the individualized neural substrates.

On the other hand, analyses based on the individual subjective responses may better capture the individual neural processes, because the neural activity and patterns generating these individualized subjective reports (behavioral outputs) may still map to brain locations in a consistent and meaningful way. If the neural processes found this way are robust and consistent enough, individual-level replication can be achieved (Smith and Little, 2018). Adopting this logic, our subsequent analyses looked at the mapping of subjective reports to brain activity, and accordingly focuses on individual-level data and results.

### The perceived action and emotion representations were high-dimensional

To investigate perceived action and perceived emotion, we examined the high dimensional space of all subjective responses across participants using principal component analysis (PCA), since all 400 responses were in the same 300-dimensional space.

The first few principal components (PCs) for the perceived action and perceived emotion representations captured a large amount of variance (First 10 PCs: 86.17% and 89.63% variance; first 50 PCs: 97.76% and 99.18% variance, **Fig. 1F, G**, analysis performed on 1^st^ to 50^th^ PC for both representations).

The perceived action representation showed considerable individual variability when compared to the 10-action-category model (**Fig. 1Ea, Ga**). The first and biggest few components did not show clear separation of action categories. Instead, individual variability was more prominent (**Fig. 1H**, correlation to individual subject model in the range of 0.1 to 0.2), while it remains elusive what accounts for the majority of the variance in PC 1. Only the later and much smaller PCs were correlated more to the 10-action-category model (7^th^, 9^th^, 10^th^ PCs), the non-emotion/emotion model (7^th^, 22^nd^, 6^th^ PCs), and the implied motion ratings (6^th^, 10^th^, 18^th^ PCs), and also showed relatively clear category separations (**Fig. 1H, I**). The presence of smaller PCs correlated to the action categories, emotions and implied motion ratings indicates that the perceived action representation is rich and high-dimensional. Since the categorization process could be thought as drawing a hyperplane to separate a cloud of dots in a high-dimensional space, such high-dimensional action representations potentially supports extracting the relevant categorical and continuous information, but it is not equivalent and not reducible to one of the predefined category structures.

The perceived emotion representation was lower-dimensional than the perceived action representation (79 versus 174 independent PCs), and showed higher inter-individual consistency (**Fig. 1Eb, Gb**, much lower correlation value to the individual subject model in **Fig. 1H**). The lower dimension may either be due to the fewer emotion than action categories (4 vs 10), the higher inter-individual consistency, or is an intrinsic property of the emotion representation. The first few PCs showed a moderate correlation to the non-emotion/emotion model and valence ratings, and even to the 10-action-category model (**Fig. 1H, J**). Interestingly, visual inspection of the RDMs showed that the perceived emotion representation contained the separation of non-emotion and emotion categories (**Fig. 1Eb**, PC1 and 2 in **Fig. 1Gb**), but also contained finer-grained separation for individual emotions. These were captured by the 3^rd^, 4^th^ and 6^th^ PCs, corresponding to happy, fearful and sad (**Fig. 1Gb**, see the locations of the prominent white/red bars across all conditions, showing similarity within the category and dissimilarity to the other 9 categories). In the 3^rd^ and 4^th^ PC, the angry emotion could further be separated. This indicates that the perceived emotion representation is also a high-dimensional one; brain areas containing such representations could potentially support both the non-emotion/emotion categorization, and categorization of each individual emotions.

### Univariate results were consistent with the literature

For the fMRI data, as a sanity check, we first performed conventional group-level univariate analysis with data smoothing, for the 10 action categories, and parametric modulations for implied motion and valence (**Figure 2**). The 7T data were very robust: the activation location and time course profiles at 7T were similar to 3T ones, but showed higher %-signal change (See **Figure S2**). Participants remained attentive to the stimuli (mean response accuracy for catch trials was 92.49%, SD=5.44%, the main mistakes being misses and false alarms, number of error trials ≤ 3 per participant).

**Figure 2.**
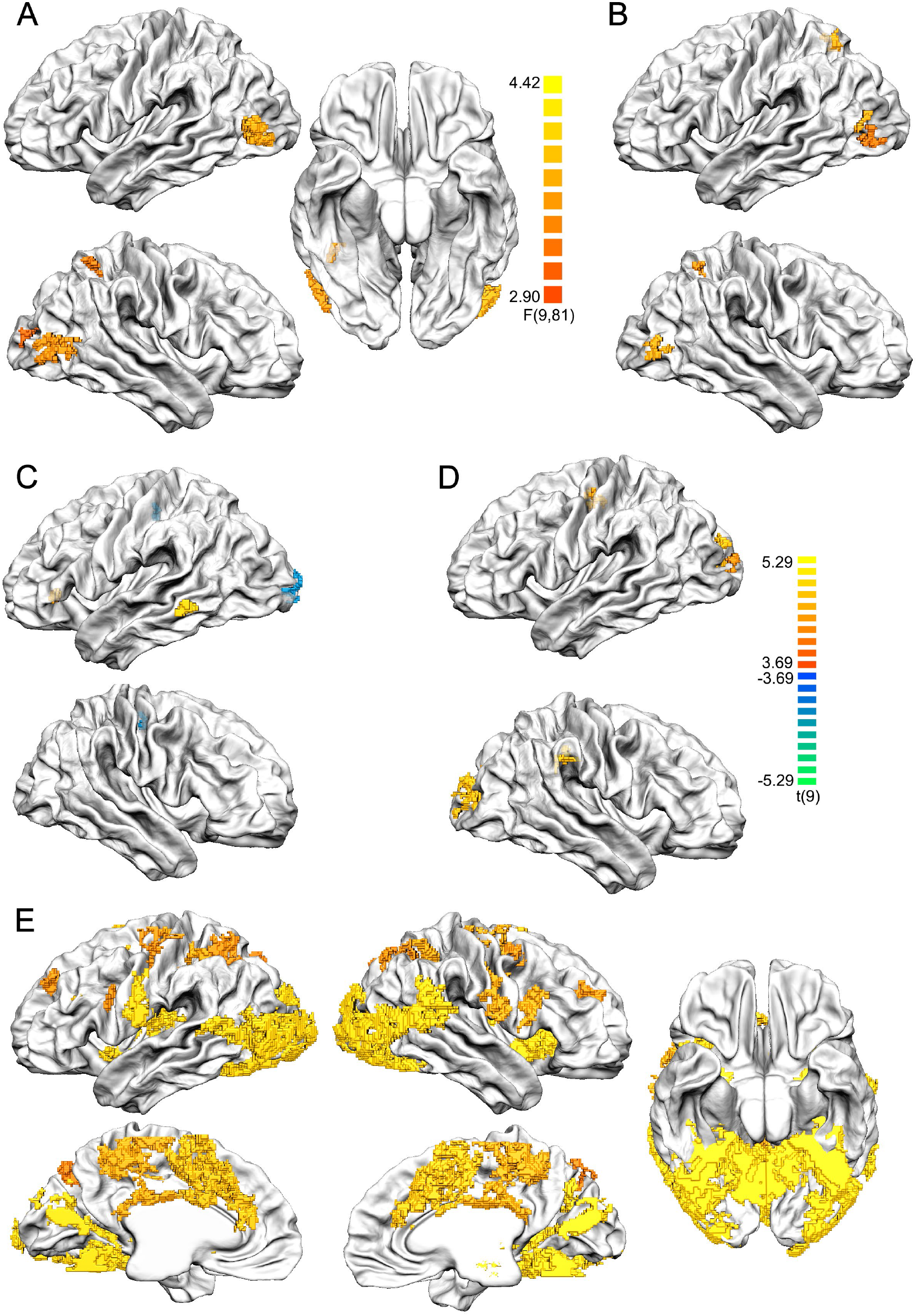
Univariate results. Data in volume space were plotted on the group-aligned surface mesh after cortex-based alignment. Functional data were smoothed at 6 mm FWHM, all maps were cluster-size thresholded with Monte-Carlo simulation, alpha=0.05, n simulations=5000. We used the initial p<0.005 to alleviate the false negatives due to high anatomical inter-individual variability, which was clearly observed in our functional localizer data. **A**. ANOVA of 10 categories. Color bar p range: 0.005 to 0.0001. **B**. Parametric modulation of rated implied motion. **C**. Emotional categories > non-emotional categories (excluding neutral standing still). **D**. Parametric modulation of rated valence. **E**. 10 categories > baseline. Color bar p range in **B to E**: 0.005 to 0.0005. Cluster size thresholds in **A** to **E**: 115, 55, 46, 56, 128 functional voxels.

The 10-category one-way ANOVA showed differences across action categories in bilateral EBA, right FBA, right lateral occipital cortex (LOC) and right mIPS (**Fig. 2A**). Similarly, the parametric modulation of implied motion showed higher activity for actions rated with higher implied motion, in bilateral EBA, right mIPS, left pIPS, right rostral cingulate zone (RCZ), where the former 3 overlapped with the ANOVA clusters (**Fig. 2B**). The 10 categories > baseline contrast showed activation in body-sensitive areas, and areas of the action observation network (**Fig. 2E**), consistent with the literature (see de Gelder and Poyo Solanas, 2021 for a review).

When contrasting emotional and non-emotional categories (excluding the standing-still condition), the left IFG and left middle temporal gyrus (MTG) showed higher activity for emotional categories, while early visual areas (EVC) and right precentral gyrus showed higher activity for non-emotional categories (**Fig. 2C**). The activation map was very similar when including the standing-still condition. These univariate results were consistent with previous findings (de Gelder et al., 2004; Dricu and Frühholz, 2016; Kober et al., 2008; Molenberghs et al., 2012; Sinke et al., 2010). However, for the parametric modulation of valence, we only observed clusters modulated by positive valence, but not for ratings of negative valence (negative modulation). These clusters were found outside the frontal lobe, in bilateral early visual cortices (including bilateral V3a and right calcarine sulcus/cuneus), right posterior collateral sulcus, right supramarginal gyrus, left precentral gyrus/central sulcus (**Fig. 2D**).

We cannot exclude that some of these may be false negatives due to the exacerbated inter-individual variability specifically at 7T, where activation clusters were small, and were constrained to the gray matter even after spatial smoothing. This effect could be observed in the functional localizer data: despite that we found robust EBA, FBA, FFA clusters in all individual participants (data smoothed 3 mm FWHM, contrasts: bodies>faces, houses, tools, words; faces>bodies, houses, tools, words), and the activation sites were reliable in one participant between 3T and 7T scans on different days (**Figure S2**), the across-participant overlapping of these clusters were low; the group-level GLM (data smoothed 6 mm FWHM) only showed an R EBA/pSTS cluster for bodies, L EVC and bilateral precuneus clusters for faces (**Figure S3**). Thus for whole-brain activation and searchlight analyses, we used the initial p threshold of 0.005 to balance between the false positives and false negatives.

### Predefined non-emotion/emotion and 10-action-category representations in the brain

Next we followed the conventional predefined-model-driven approach, and performed RSA searchlight analysis for the RDMs of the 10 action categories, and the non-emotion/emotion categories. We constructed neural RDMs from univariate t maps of the 40 individual stimuli (no smoothing of functional data, Pearson’s correlation distance), and performed RSA searchlight (radius=5 voxels; Spearman correlation to the predefined category RDMs, one-sampled t test against 0 for the Fisher’s Z-transformed rho values at the group-level, cluster size thresholded at alpha of 0.05, Monte-Carlo simulation n=5000). The same group-level test and cluster thresholding scheme was used for all searchlight analysis throughout the study.

For the predefined non-emotion/emotion categories, we found two areas positively correlated to the model RDM, in L central sulcus (adjacent to the cluster found in parametric modulation for valence ratings in the univariate analysis, in **Fig. 2D**), and R PMd/FEF (**Figure 3B**). Another four areas showed negative correlation to the RDM, in L precuneus, L caudal cingulate zone, bilateral medial SFG, and L thalamus. These negative correlations may indicate fine-grained processing of individual stimulus exemplars within each category (here the non-emotion or emotion categories), or similar processing across categories. Since their interpretation would require further systematic examination, we limit our interpretations on positive correlations.

**Figure 3.**
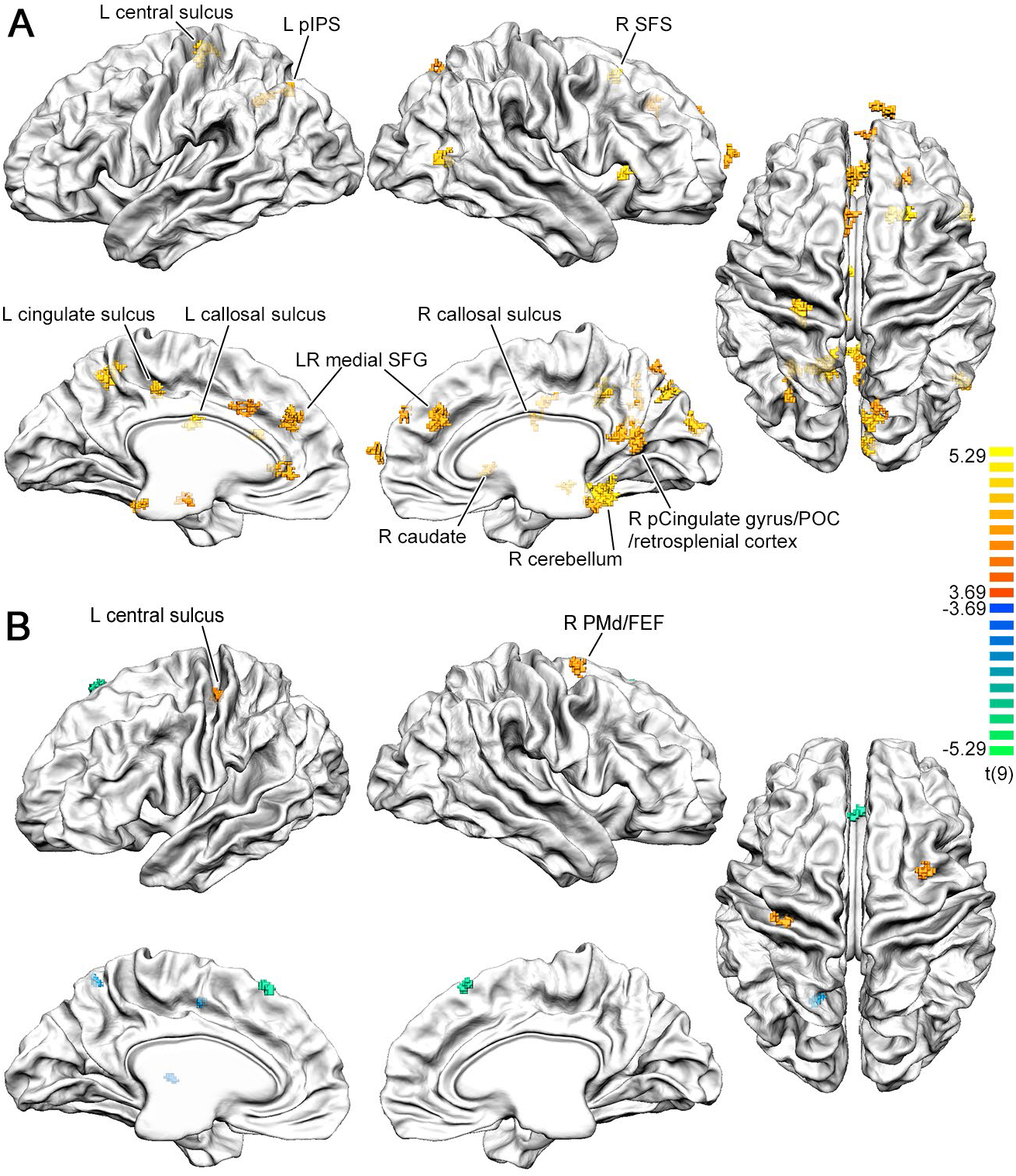
Searchlight RSA showed multiple areas correlated to the predefined category models. **A**. Results for the 10-action-category RDM, cluster size threshold=43 voxels. The labeled areas were ones overlapped with the perceived action and emotion FC networks. **B**. Results for the non-emotion/emotion RDM, cluster size threshold=35 voxels. The areas labeled were ones with positive correlations to the RDM. Color bar p range in both **A** and **B**: 0.005 to 0.0005. Cluster size thresholds: 43, 35 functional voxels. Initial p<0.005.

For the predefined action categories, we found 29 brain areas positively correlated to the 10-action-category RDM (**Figure 3A, S4, Table S3**, denoted “10-action-category areas” below), a lot of which were medial areas, and no negatively-correlated areas. When thresholding at p<0.001, eight of these areas were still present, including the R cuneus, R EBA/hMT+, L central sulcus, L precuneus, R posterior cingulate gyrus and sulcus, bilateral medial superior frontal gyrus, anterior RCZ. Some of these 10-action-category areas consistently showed group-level univariate activation for multiple action categories, in body or action perception-related areas including R EBA/hMT+, L pIPS and L RCZ; but several areas did not show above-baseline univariate activation for any action category, including L central sulcus, R anterior SFS, R cerebellum, L vmPFC and bilateral medial SFG (**Figure S5, Table S4**).

We next examined the task-residual functional connectivity, which presumably reflects information transfer, and displays certain levels of consistency with structural connectivity (Bullmore and Sporns, 2009), and thus could potentially capture the direct upstream/downstream areas for a seed area. Using these 29 areas as seed regions, we observed fine-grained connectivity patterns for each seed region, both showing hemispheric symmetry. These areas were highly interconnected, and were also connected to the action-observation-related areas (EBA, FBA, IPS, pSTS, SMA, PMd, PMv, M1, cerebellum), and DMN areas (dmPFC, vmPFC, precuneus, TPJ, mSTS). Furthermore, most areas were heavily connected to bilateral caudate, putamen and thalamus; and 14 out of 29 areas were connected to the hippocampus. See supplementary **Figure S6** for a summary of these connectivity patterns.

### The representations of low- or mid-level visual features and higher-level attributes

An important question to disentangle is, whether the brain areas showing category boundaries contained representations for the more abstract categories per se, or merely for the mid-level and/or lower-level visual feature differences between the stimuli, which could co-vary with the abstract category boundaries.

To investigate whether some of the 29 brain areas found by the 10-action-category RDM may be related to some low/mid-level visual features, we performed RSA regression, using the RDMs of higher-level stimuli attributes and low- or mid-level visual features as predictors. We only tested a limited number of features, motivated by the literature as in our previous studies (Poyo Solanas et al., 2020a, 2020b), because the model space is infinite and impossible to cover by our current study. High level stimulus attributes included non-emotion/emotion, implied motion, valence, and actor identity. The low level features included the raw pixel values of each stimulus picture, and the body joint coordinates, extracted from each stimulus picture by the OpenPose library (Cao et al., 2019). The presumed mid-level visual features were computed from the body joint coordinates,, including features related to body-part (head, shoulder, waist) orientation, as these may signal the directions of social interactions; the hands/feet-to-head distances, which may be related to the peripersonal space (Bufacchi et al., 2016); and the relaxation of the arms/legs. All RDMs were put in the same linear model, so that the unique contribution of each RDM could be examined.

Most of the low-mid-level RDMs indeed correlated to the 10-action-category RDM (rho ranging from 0.08 to 0.36), except the raw-pixel-value and head-orientation RDMs. See **Figure 4A, B**. These correlations are to be expected, because the actors of the stimuli set received specific instructions per action category (Stienen and de Gelder, 2011) (see all stimuli in **Figure S1**).

**Figure 4.**
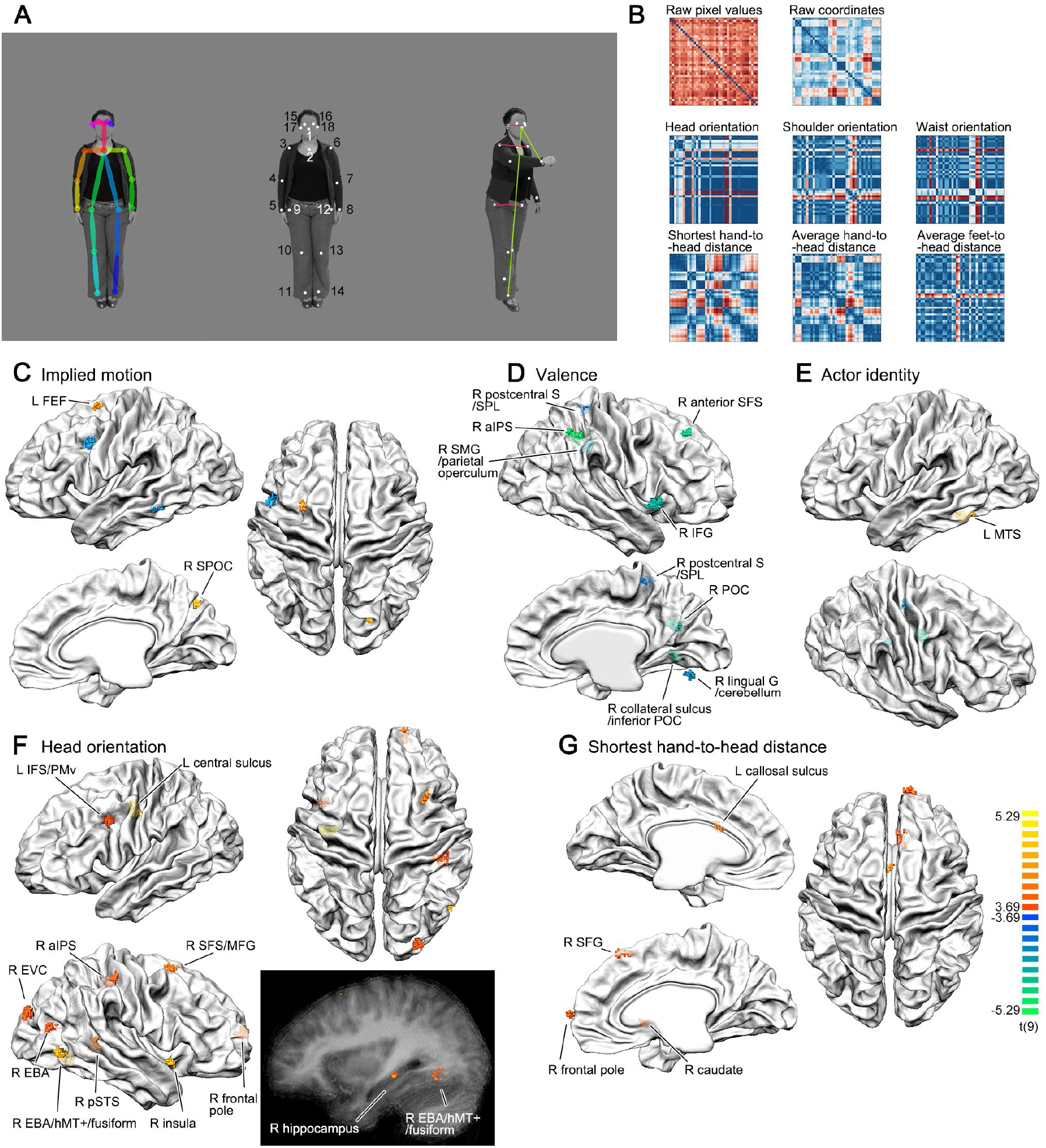
Low/mid-level features (**A** and **B**), and whole-brain searchlight results corresponding to these features (**C** to **G**). **A**. Skeleton in the left panel: coordinates of the 18 joints extracted by the OpenPose library; white dots in the middle panel: joint coordinates after manual adjustment; right panel, magenta lines: distances used to compute head, shoulder, waist orientations, normalized by the corresponding distances in the neutral stand-still stimulus of the same actor (see middle panel); green lines: hand/foot-to-head distance. **B**. The low/mid-level features, showing RDMs for stimuli set A. The raw coordinates RDMs were computed excluding joints on the ears and eyes, but was very similar to the ones computed including these joints (RDM similarity rho=0.976, 0.98 for stimuli set A and B). Most of these RDMs correlated to the 10-action-category RDM. **C, D, E**. Searchlight results for higher-level attributes. The valence RDM did not show positively correlated clusters. Cluster size thresholds: 38, 44, 31 functional voxels. **F, G**. Searchlight results for mid-level features that showed clusters positively correlated to the corresponding RDMs. Cluster size thresholds: 48, 37 functional voxels. Color bar p range in **C** to **G**: 0.005 to 0.0005. Initial p<0.005. Clusters showing positive correlations to the corresponding RDMs were labeled. In **D**, clusters showing negative correlations were also labeled.

We found that some of these stimulus features and attributes could partially explain the activity in some of the 10-action-category brain areas (with a positive beta estimate significantly bigger than 0). For the low-level features, the raw pixel values representation could be found in R cuneus and R CCZ (negative beta in R SPOC); the raw-joint-coordinates representation could be found in R EBA (negative beta in R callosal sulcus). For mid-level features, the head orientation representation could be found in R SPOC (negative beta in R dmPFC); the shoulder orientation representation was found in L pIPS, and the shortest hand-head distance representation in R cerebellum. For higher-level attributes, only the actor identity representation showed a positive beta estimate again in the R SPOC (negative beta in R cuneus).

We then performed whole-brain searchlight analysis for all those attributes and features (**Figure 4 C** to **G**). For high-level features, we did not find areas positively correlated to the valence rating RDMs. The implied motion rating RDMs showed positive clusters in L FEF (adjacent but not overlapping with the L PMd cluster for perceived emotion) and R SPOC. And interestingly, the actor identity RDM showed a positive cluster in L MTS, close to EBA and FBA.

We found positive clusters for two mid-level visual features: head orientation and shortest hand-to-head distance, but not for any other low- or mid-level visual features, despite their similarity to the two features with positive clusters (shoulder and waist orientation RDMs were correlated to the head orientation RDMs: rho ranges from 0.317 to 0.412; the shortest hand-to-head distance RDMs were correlated to the average hand-to-head distance RDMs, rho=0.593, 0.646 for stimuli set A and B). This indicated that these two features may be biologically meaningful ones for the brain.

### RSA searchlight analysis revealed representations for perceived action in left mSTS and for perceived emotion in left PMd

We next returned to the individual subjective reports, searching for similar neural representations for the individualized perceived action and emotion RDMs. We found one positively correlated cluster for each of the subjective report RDMs, in left mSTS for perceived action, and in left PMd for perceived emotion, indicating that these two areas may be involved in the subjective understanding and reporting of action and emotion, respectively. The L PMd cluster is consistent with the analysis based on predefined non-emotion/emotion representation, where the R PMd/FEF cluster was at a very similar location on the other hemisphere to the PMd cluster found here. All other clusters were negatively correlated to the subjective category RDMs. See **Figure 5** and **TableS5** for the complete list of clusters.

**Figure 5.**
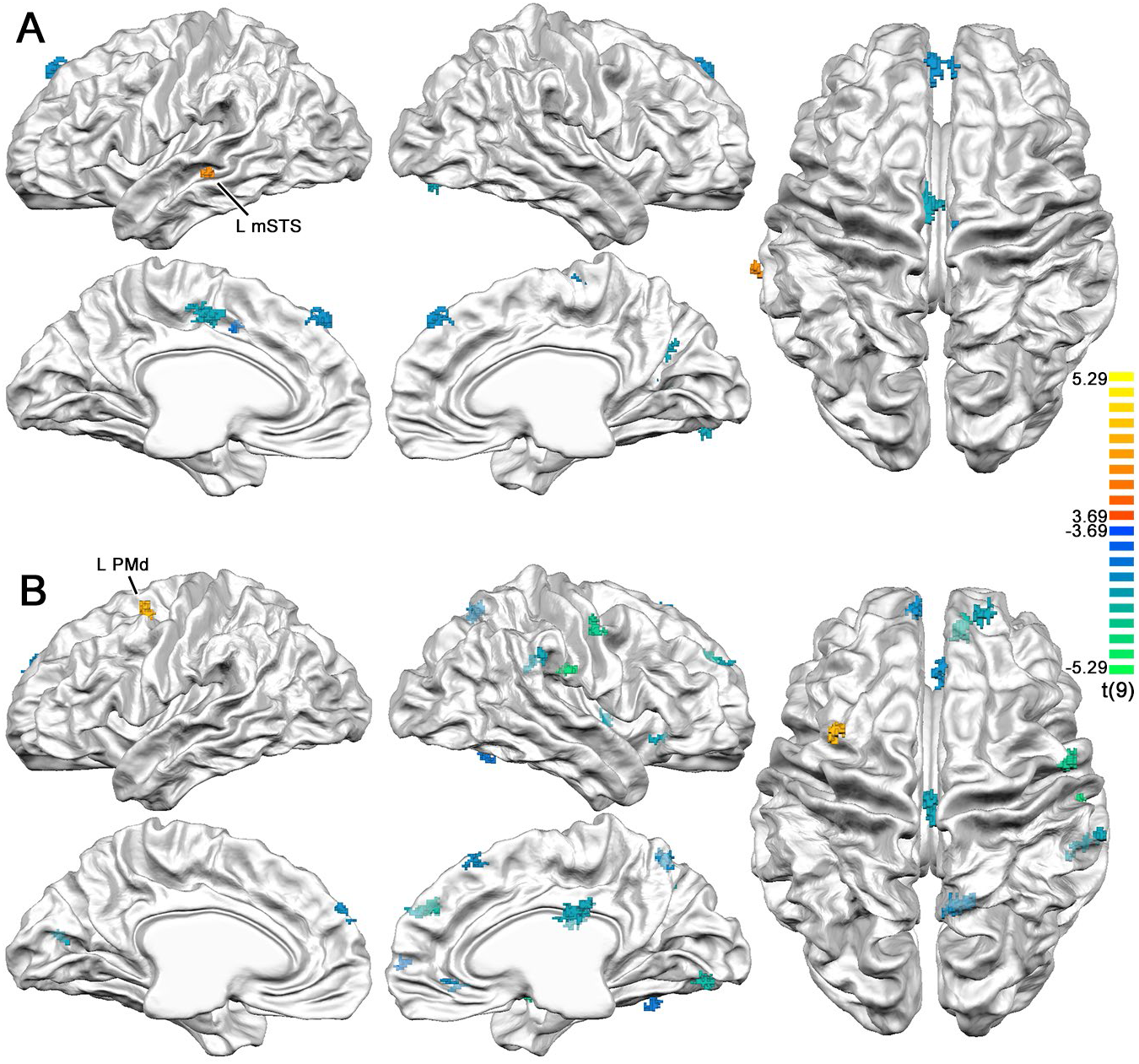
Searchlight RSA revealed corresponding clusters to the perceived action model in L mSTS **(A)**, and to the perceived emotion model in L PMd **(B)**. Color bar p range: 0.005 to 0.0005. Cluster size thresholds in **A** and **B**: 39, 38 functional voxels. Initial p<0.005.

Interestingly, the L mSTS and L PMd clusters did not show reliable univariate activation for the 10 action categories (one-sample t test of percent signal change against baseline, all p>0.052, except in L PMd for the Anger category, p=0.031), thus they could not be found by conventional univariate analysis. Neither was the 10-category univariate ANOVA significant in these two clusters, p=0.309 and p=0.945 respectively.

### The putative direct upstream/downstream areas of L mSTS and L PMd identified by task residual functional connectivity and hierarchical clustering

The L mSTS and L PMd clusters were found by their representation similarity to individualized, subjectively reported actions and emotions, which already showed considerable individual variability and hard-to-interpret principle components. Thus, to further understand the involvement of these two areas in the current task, we could no longer use the standard model-comparison methodology, where a single predefined-model is compared to the neural RDMs across all participants. Instead, we used a data-driven approach based on two assumptions: (1) the information transfer stages of a given area can potentially be captured by functional connectivity; (2) if the object representations are successively re-represented and “untangled” in the ventral visual pathway (DiCarlo et al., 2012; DiCarlo and Cox, 2007), similarly the neural representations of bodily action and emotion representations are gradually transformed across a chain or a network of brain areas, from the representation of the visual input to the representation of the subjective output, without sudden changes between two adjacent stages in the information-transfer chain. In this way, we linked the functional connectivity and representational analyses together (Ju and Bassett, 2020). To find the set of brain areas in the same information-transformation chain, we performed task-residual functional connectivity to the L mSTS seed and the L PMd seed, respectively. Using hierarchical clustering in the two resulting networks, we could then find areas with neural representations most similar to the two seeds, some of which may correspond to their direct upstream/downstream areas.

#### Task-residual functional connectivity

We regressed out the individual stimuli and head motion parameters from the functional data (deconvolution, data smoothed at 3 mm FWHM), obtained task-residual time courses, and performed functional connectivity (FC) analysis (Pearson’s correlation), with L mSTS and L PMd as seed regions. See **Figure 6A, B, left panels**. This analysis revealed two partially overlapping networks for action/emotion understanding at the group level (**Figure 6D**), in action observation related areas including (bilateral when unspecified) IPS, M1, PMd, PMv, L IFG; and in DMN areas including TPJ, mSTS/MTG, precuneus, posterior cingulate cortex, dmPFC/vmPFC, mSFG, parietal occipital cortex (POC), retrosplenial cortex.

**Figure 6.**
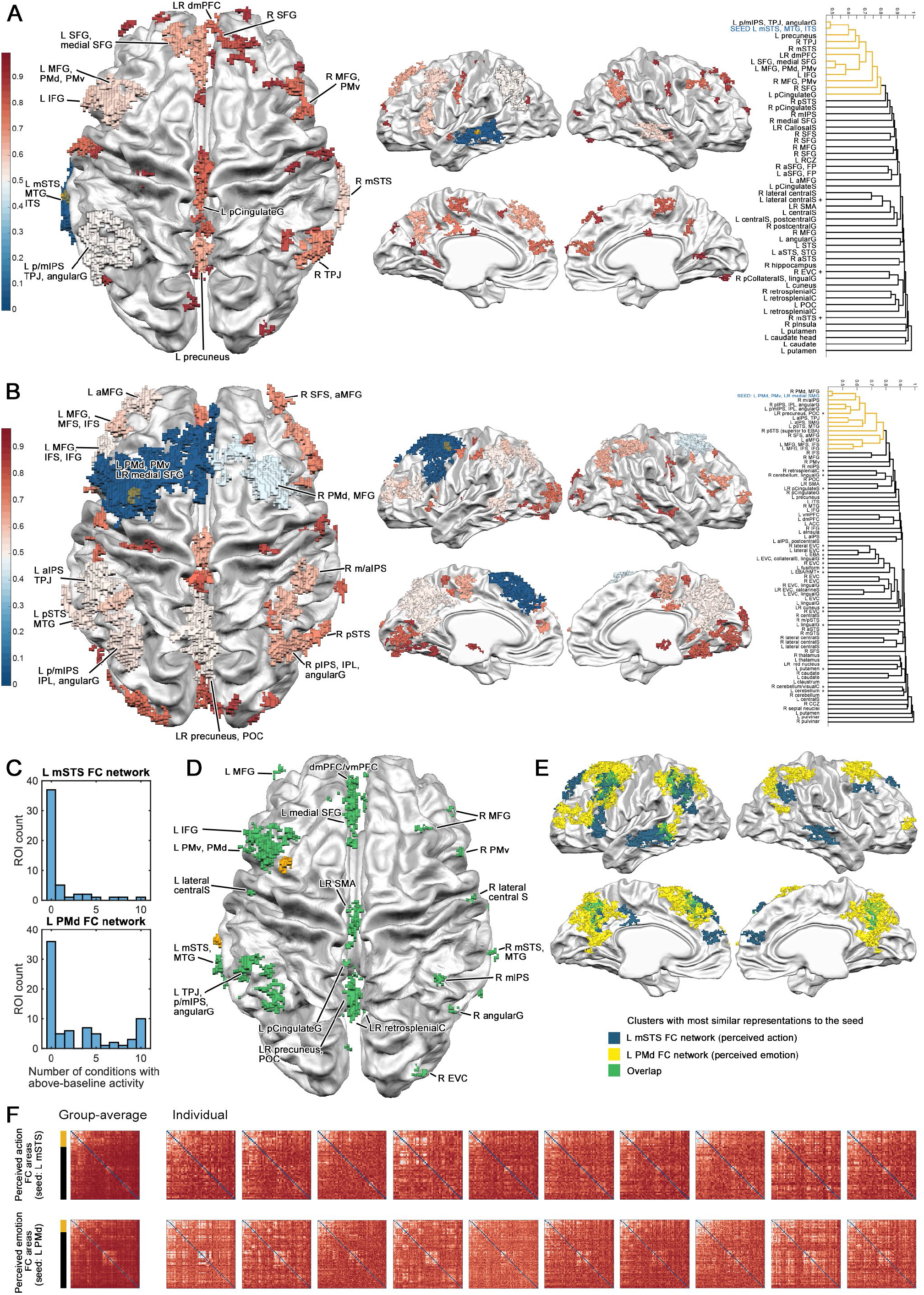
In the functional connectivity (FC) networks of the two seed regions overlapped with the DMN/semantic networks. **A** and **B**. Areas in the FC networks color-coded by pattern similarity to the seeds, L mSTS (**A**), and L PMd (**B**). Initial p<0.001, cluster size threshold in functional voxels: L mSTS network: 32; L PMd network: 40. The seed ROIs were in yellow. Color bars: representational similarities to the FC cluster containing the seed region. For each network, the hierarchical clustering dendrograms computed on group-averaged second-level RDMs across areas (Spearman distance) were plotted in the right-most panels. The areas with the most similar representations to the FC cluster containing the seed region were colored in yellow in the dendrograms (linkage distance arbitrarily thresholded at 0.8, see a larger version in **Figure S9**), and labeled in the brain maps of the left panels. The name of the areas containing the seed regions were marked in blue. The areas with above-baseline univariate activation for more than 5 action categories were marked with a “+” sign. **C**. Histograms of number of conditions with above-baseline activity, for clusters in the two networks. Most areas did not show univariate activation for any of the 10 action categories/conditions. **D** and **E**. The two FC networks overlapped with the DMN/semantic networks. **D**. The overlap of two FC networks. **E**. The overlap of the two FC networks, only showing clusters with most similar representations to the seed (the clusters labeled in **A** and **B**). **F**. The group-averaged and individual second-level RDMs. The yellow and black bars for the group-average RDM denote the same clustering of the dendrogram in **A** and **B**. See a larger version in **Figure S9**.

Interestingly, some overlapping areas in these FC networks are part of the semantic network, including mSTS, IFG, TPJ/angular gyrus, dmPFC, vmPFC, posterior cingulate cortex, retrosplenial cortex (Binder et al., 2009); and the L mSTS FC network corresponded to the semantic network especially well. See **Figure 6D** and **E**. The seed cluster L mSTS itself is an important area in the semantic network, which could be activated by written word stimuli (Binder et al., 2009). Although word-specific clusters around left mSTS were found in individual participants in the functional localizer data (words>other categories), neither the seed region nor the more extended FC cluster around the seed showed consistent group-level activation for words in the functional localizer (one-sample t test against 0, L mSTS seed ROI: mean beta=0.589; t(9)=1.418; p=0.190; L mSTS FC cluster: mean beta=0.369; t(9)=1.226, p=0.251). More interestingly, although both FC networks overlapped around L mSTS, the cluster for perceived emotion was much more posterior than the one for the perceived action, which corresponded to previous findings that the STS is an heterogeneous structure with several different functions (e.g. Hein and Knight, 2008).

In addition to the cortical clusters, the FC analysis revealed bilateral caudate and putamen clusters in both FC networks (**Figure S7, S8**), which may be involved in categorization processing (Seger, 2008; Seger and Miller, 2010). The left mSTS was also functionally connected to the R hippocampus; the left PMd was also functionally connected to the bilateral thalamus, pulvinar, cerebellum, the R septal nuclei, and the bilateral red nuclei (anatomical locations clearly observable in the T2*-weighted functional images, **Figure S8**). The involvement of the hippocampus, the septal nuclei and the red nuclei in bodily action and emotion understanding were not routinely observed in previous bodily-action perception studies with univariate methods (see de Gelder and Poyo Solanas, 2021 for a review).

Furthermore, the perceived action FC network overlapped with the 10-action-category clusters in the L central sulcus, R POC/retrosplenial cortex and bilateral callosal sulci; while clusters in the perceived emotion FC network overlapped with the 10-action-category clusters in L precuneus, L pIPS, L cingulate sulcus, R SFS, R posterior cingulate/POC/retrosplenial cortex, R caudate, R cerebellum and bilateral medial SFG (labels in **Figure 3**), again showing consistency to the analysis based on predefined categories.

#### Hierarchical clustering

We then examined the possible direct upstream/downstream areas of the two seed regions, which should show the most similar neural representations to the seeds. We performed hierarchical clustering to all clusters within each FC network, on the group-averaged second-level neural RDMs across areas (Spearman correlation distance, hierarchical cluster linkage arbitrarily thresholded at 0.8). Since both seed regions were encompassed by more extensive clusters with the shortest hierarchical clustering distance to the seed (cluster in the perceived action FC network: spanning L mSTS, MTG and ITS; cluster in the perceived emotion network: an extensive one spanning L PMd, PMv and LR medial SMG), we plotted the extensive clusters (denoted as “seed clusters” below) instead of the seed regions themselves, in the dendrograms and second-level RDMs. See **Figure 6A, B, Figure S9**.

In the perceived action FC network, the FC cluster with the most similar representation (shortest distance in the dendrogram) to the seed cluster was one spanning L p/mIPS, TPJ, angular gyrus. The other areas under the same branch of the dendrogram were: the R mSTS contralateral to the seed; action-perception related areas including bilateral PMv, MFG, L PMd, L IFG; DMN/semantic areas including L precuneus, R TPJ, L posterior cingulate gyrus, LR dmPFC, LR and medial SFG.

In the perceived emotion FC network, the FC clusters with the most similar representation to the seed cluster was one spanning R PMd and MFG, contralateral to the seed. The other FC clusters in the same branch were: L pSTS, MTG, R pSTS, action-perception related areas including bilateral IPS, IPL, angular gyrus, L IFS, IFG; DMN/semantic areas including L TPJ, bilateral precuneus, POC; also the lateral and anterior prefrontal cortices including bilateral MFG, anterior MFG, L MFS, R SFS.

These clusters may be directly involved in understanding the bodily actions and emotions. In both FC networks, it is interesting to observe the strong involvement of both action-perception related areas and DMN/semantic areas, showing symmetry across hemispheres. Especially for the DMN/semantic areas, their direct involvement in bodily action/emotion understanding and interactions with the action-perception related areas were not reported in previous studies. This may be due to the fact that most of the areas in the two FC networks did not show an activation level different from baseline (one-sample t-test of beta values per action condition against baseline, FDR corrected across the 10 action categories per ROI). See **Figure 6A, B, C**. Thus, they were unlikely to be reported when using only univariate methods.

The clustering in individual participants was relatively consistent, with moderately correlated second-level matrices across participants (averaged Spearman Rho, perceived action: 0.418, SD=0.0473; perceived emotion: 0.374, SD=0.0578). Also, the second-level pattern-similarity clustering structure across areas does not seem to be linked to the low-level visual features from the stimuli, but seem to be a real organization feature, because the clustering structure could be seen in all 10 participants, despite the fact that each 5 participant saw a different stimuli set (set A and B). See **Figure 6F**.

Apart from the clustering showing high similarity to the seed clusters in every participant (upper-left side of each RDM), we saw another prominent clustering of areas in each of the two FC networks of individual participants (27-32^nd^ areas in the perceived action FC RDM, 36-51^st^ areas in the perceived emotion FC RDM, both in the middle of the RDMs, see an enlarged version in **Figure S9**). However they showed lower similarity to the two seed regions (red shade for FC clusters in **Figure 6A, B, left panels**), indicating that these areas are at different processing stages than the final output stage of the seed regions. In these two groups of clusters, despite their very different univariate activation levels, their multivariate RDM patterns were clustered in the same branch of the dendrogram. The clustering in the perceived action FC network consisted of areas around the bilateral central gyrus (lateral-central/central sulcus, post-central gyrus, bilateral SMA), most showing above-baseline activation for 0 to 3 of the 10 action categories, apart from the L lateral central sulcus showing activation for 8 categories. The clustering in the perceived emotion FC network consisted of areas in the bilateral ventral-lateral visual pathway (EVC, lingual gyrus, collateral sulcus, fusiform, EBA/hMT+, lingual gyrus, cuneus), where 8 areas showed above-baseline activation for all 10 action categories, another area showing activation for 8 categories, the other 7 areas showing no activation for any of the categories.

Combining the results of functional connectivity and hierarchical clustering, our two assumptions of the information-transfer-chains seems to be able to discover meaningful and replicable functional organizations in the brain.

## Discussion

Our goal in this 7T fMRI study was to go beyond the analysis based on predefined action and emotion categories, and examine how participants perceived bodily actions by analyzing participants’ subjective understanding of the actions and emotions displayed. Dimension reduction (PCA) revealed that subjectively perceived action and emotion representations were high-dimensional and could not be reduced to the predefined category representations, despite being correlated to them in smaller principle components. Some emotional categories were reflected in the smaller principle components of the perceived emotion representation. Clusters in L mSTS and L PMd had representations that were the most similar to the perceived action and emotion representations. Areas located in the action-observation network and the semantic network/DMN were functionally connected to these two clusters and also showed similar multivariate patterns. This provided direct evidence for the involvement and interplay of both networks in transforming the same visual inputs into individualized subjective understanding of bodily actions and emotions.

For the predefined categories, the non-emotion/emotion representation corresponded well to the perceived emotion representation both behaviorally and neurally, showing a cluster in R PMd/FEF, contralateral to the cluster for perceived emotion. For the predefined 10-action-category representation though, the resulting clusters were more numerous, and possibly confounded by low/mid-level visual features that may co-vary with action/emotion categories. Although less clear than the subjective report analysis, all of the clusters were strongly connected to both the action-observation network and the semantic network/DMN, consistent with the perceived action results. For mid-level visual features, we further found that the head orientation and the shortest hand-to-head distance were represented in the brain, indicating that these two features may be important for body perception.

Comparing the analysis of predefined categories with that of the subjective reports, our results suggest that both analysis would yield similar results in the brain, when the subjective reports are moderately correlated to the predefined categories (rho=0.491 to the non-emotion/emotion RDM) and have moderate individual variability. However the results would converge less when the subjective reports are less correlated to the predefined categories and have high individual variability. In this case the individualized analysis of subjective reports would be more precise, and capture additional information missed by predefined models, or group-averaged individual models. The convergence of these two kinds of analyses needs further study.

### The involvement of the DMN/semantic network in both action and emotion understanding

At first sight, it may seem that the involvement of the DMN/semantic network found in the current study simply results from the use of word embeddings. We argue that this is not the case, and that our study reveals the individual brain underpinnings of action and emotion understanding as involving both the action observation network and the DMN/semantic network. Although the word embeddings indeed represent the semantic distances between the words, these semantic distances reflect the similarities between different concepts. Our perceived action and emotion representations were different, despite being computed from word vectors from the same 300-dimensional space. Also, the perceived emotion cluster in L PMd is not a key node area in the semantic but rather in the action observation network, but nevertheless was revealed being connected to the DMN/semantic network.

Actually, the involvement of IFG (Caspers et al., 2010; de Gelder et al., 2004; Dricu and Frühholz, 2016; Molenberghs et al., 2012) and areas in the DMN/semantic network (Chikazoe et al., 2014; Peelen et al., 2010; Skerry and Saxe, 2015) has been consistently found in action perception and emotional expression perception, although in separate studies and in different contexts. For the DMN and especially for the vmPFC area, it was related to abstract representation of emotional stimuli found with multivariate RSA (Chikazoe et al., 2014; Peelen et al., 2010; Skerry and Saxe, 2015), rather than with simple univariate contrasts. The simultaneous involvement of the IFG and the DMN in action understanding was found in one univariate study contrasting participant’s attention on either the intention or means of the performed actions (de Lange et al., 2008). In that study, the IFG activity was higher for actions with unusual intentions compared to usual ones; the DMN was instead showing higher activity when participants were paying attention to the intentions rather than the means of the action.

Two recent studies provided more evidence consistent with ours. One RSA fMRI study of observed actions (Tucciarelli et al., 2019) used a large set of action categories, constructed a semantic similarity model of the meanings of the actions as well as other similarity models obtained from individual participants’ behavioral sortings of similarities, and searched for corresponding RDMs in the brain. They found bilateral clusters in IFG/PMv, pIPS, and lateral occipitotemporal cortices (LOTC, close to EBA) corresponding to the semantic similarity model, although only the left LOTC cluster remained after RSA regression controlling the effects of other models. That study controlled the inter-individual consistency of the stimuli perception during the stimuli selection process, which was not done in the current study, and may explain the discrepancy. Another study (Lee Masson and Isik, 2021) examined the fMRI responses during naturalistic movie watching using encoding models, and found that a social-affective model (including features of an agent speaking, social interactions, theory of mind, perceived valence and arousal) significantly explained the fMRI response in the left STS across two different movies. This cluster fully overlapped with our L mSTS cluster.

In fact, the IFG and the areas in the DMN were all parts of the semantic network, thus their involvements could also be studied in the future in the context of the semantic network, apart from the context of “mentalizing” usually associated with the DMN. The importance of the semantic network in emotion understanding is further supported by behavioral studies of semantic dementia patients. In one such studies, three semantic dementia patients with left temporal pole atrophy and impaired semantic knowledge were asked to sort faces into piles by emotion. These patients were not able to distinguish between emotional faces with negative valence, despite that their perception of positive/negative affect, the visual features for each facial emotion and identity were intact (Lindquist et al., 2014).

Although we did not examine the temporal pole due to coverage limitation, our multivariate RSA and functional connectivity results support the importance of the DMN/semantic network in action and emotion understanding, and are consistent with the recent view that the DMN is potentially the network that integrates extrinsic and intrinsic information (Yeshurun et al., 2021). Our results are also consistent with the literature, which reported that the DMN did not show above-baseline activation for passive action observation without deliberating the goals/intentions (Van Overwalle and Baetens, 2009). We further revealed that both the DMN/semantic network and the action observation network were involved in the process of action and emotion understanding, that they were consistently found in all 10 participants within the same functional connectivity networks and showed similar multivariate patterns despite very different levels of univariate activation. The multivariate methods worked in our study and in previous RSA studies, because they took the multiple dimensions in the high-dimensional neural data into account, while the univariate method considers only one or a few specific dimensions which associated with specific contrasts (Haxby et al., 2011). Our study stresses the importance to further examine the function of the DMN/semantic network in action and emotion understanding with multivariate methods in future studies.

### Understanding versus categorization

Previous experiments have mostly used explicit emotion and action categorization tasks with predefined categories, and implicitly used explicit categorization as a proxy for subjective understanding. However, categorization and understanding may involve different neural substrates, as categorization involves some level of abstraction. Our study could not differentiate between these two processes, because first, the RSA method could not disentangle categorical boundaries driven by lower-level visual features, or concrete perceptual categories (Hoemann et al., 2020; Mansouri et al., 2020) that were bound to individual exemplars, or abstract categories that generalize across exemplars. Second, what we obtained in the subjective reports were descriptions or labels of action and emotion for each individual stimulus, rather than more abstract categorization to the group of stimuli. Thus, the resulting L mSTS and L PMd clusters for perceived action and emotion may not represent the most abstract level, and the more abstract categorization may happen in other areas functionally connected to these two areas, and could perhaps utilize these category boundaries in computation. The different levels of abstractions may also have driven the discrepancy between the analyses with subjective reports and predefined categories, although both analyses pointed to the strong involvement of the caudate, which is in the executive loop of categorization learning tasks (Seger, 2008; Seger and Miller, 2010). The areas found in our study could serve as target areas in future studies, to examine the level of abstraction.

### Brain areas for emotion understanding

Our analysis with the predefined non-emotion/emotion category model and with the perceived emotion model pointed to the PMd areas in the right and left hemispheres. This area may contain information exhibiting categorical boundaries between non-emotional/emotional stimuli, but again the level of abstraction could not be disentangled. Future studies could use emotional facial or voice stimuli, accompanied by subjective reports, to examine whether this area is specific to bodies and actions, or is more general for emotions in different modalities (Vaessen et al., 2019b).

We did not find evidence for coding of valence in frontal (higher-order) areas, either with monotonic univariate activity modulation or with multivariate representations. However, valence coding may be bivalent, such that the vmPFC/mOFC activity monotonically increased for both positive and negative valence, as found in an RSA study (Chikazoe et al., 2014). This might be the case, as we found evidence that the vmPFC was potentially showing a fine-grained pattern between similarly perceived emotions (**Figure 5B**). With only one category of positive emotion, we were not able to examine the possible bivalent activity.

In the analysis of the predefined action and emotion models, and in the univariate parametric modulation of valence rating, we found adjacent/overlapping clusters in the L central sulcus (**Fig. 2D, Fig. 3**). These clusters are also adjacent/overlapping to the two FC networks for perceived action and emotion (**Fig. 6AB**), but with different neural representations to the two seed regions for perceived action and emotion. This indicated that the primary sensorimotor areas were involved in action and emotion processing, although the representation there may be of lower-level features and may not be emotion-specific.

### Importance of using individualized subjective reports to study higher-level cognition

With our subjective-report-based analysis, we found that the perceived action and emotion representations were more high-dimensional and multi-faceted than the predefined category representations, consistent with the recent series of studies examining subjective reports (Cowen et al., 2019; Cowen and Keltner, 2021, 2020, 2017). We also found neural representations corresponding to these subjective report representations in higher-level areas outside the ventral and dorsal pathways.

Apart from a few RSA studies which linked individualized behavioral data to the brain data (Chikazoe et al., 2014; Stolier and Freeman, 2016; Tucciarelli et al., 2019), many previous RSA studies either utilized only the predefined categories (Peelen et al., 2010), or the behavioral ratings from an independent group (Bracci and Op de Beeck, 2016; Mur et al., 2013; Peelen et al., 2014; Skerry and Saxe, 2015; Vaessen et al., 2019a), or the averaged behavioral responses of the same participants scanned (Connolly et al., 2012; King et al., 2019). For high-level cognition that has a more remote relation with the sensory stimuli and has more variability between individual participants, the use of individualized behavioral data may be of particular interest, as shown for object recognition (Charest et al., 2014), traditionally thought to have considerable behavioral judgment similarities between participants. In precision fMRI that densely samples individual participants, it has also been found that a large part of the variance in the functional network similarity was explained by factors related to the individual subjects (showing high functional network similarity within individual), more in top-down control systems than sensorimotor systems (Gratton et al., 2018). Such individual network variability may partially be accounted for by subjective experience, which could be objectively quantified by multiple easy-to-use pre-trained word embeddings.

#### Advantages and limitations of the current 7T experiment

The use of high-resolution 7T fMRI in the current study has advantages and limitations. With higher gray-white-matter contrast and higher temporal signal-to-noise ratio, the resulting data were very robust (see the functional localizer data in **Figure S2 and S3**), the activation clusters in individual participants were highly localized in the gray matter, and we observed small subcortical clusters in multiple analyses at the group level, including periaqueductal gray, medial geniculate nucleus, substantia nigra, red nucleus, and the septal nuclei. However, the current 7T data also induced the possibility of false negatives, where the inter-individual anatomical/functional variability was exacerbated and would no longer be compensated by extensive smoothing, as observed again in the functional localizer data (**Figure S3**). Better whole-brain group-level analysis schemes apart from the ROI analysis are needed to benefit from both the high functional resolution and the large brain coverage, which potentially include more fine-grained functional parcellations (Glasser et al., 2016; Yeo et al., 2011) and hyperalignment (Haxby et al., 2011).

## Materials and Methods

### Participants

The data of 10 healthy right-handed participants recruited from the campus of Maastricht University were included in the analyses (mean age =23.4, SD=1.955, 5 females.) Two more participants took part in the study, but due to excessive head motion observed during the scan, their scanning sessions were either aborted, or data excluded from the analyses. We planned this sample size according to previous high-resolution 7T studies, and because we perform individualized analysis while aiming replicability of effects at individual participants (Smith and Little, 2018). All participants had normal or corrected-to-normal sight and had no history of psychiatric disorders. Participants provided written consent before the study and received monetary reward afterwards. The experiment was approved by the ethical committee of Maastricht University, and was carried out in accordance with the declaration of Helsinki. The experiment was conducted in English.

### Data acquisition

The MR data were acquired in a 7T Magnetom full-body scanner (Siemens, Erlangen, Germany) in Scannexus, Maastricht University, with a Nova 1-transmitter/32-receiver head coil (Nova Medical, Wilmington, USA). Dielectric pads were used for all participants except S10 (limited by the head size), roughly covering bilateral occipito-temporal lobes. The stimuli were back-projected onto a screen behind the participants’ head (Projector: Panasonic PT-EZ57OEL, projected screen size 30 × 18 cm, resolution 1920 × 1200 pixels, refresh rate = 60 Hz, viewing distance ∼99 cm, screen visual angle 17.23 × 10.38 degrees) and the participants viewed the screen through a mirror fixed on the head coil. Participants came for two scanning sessions, a 2-hour session for the main experiment, and a 1-hour session for functional localizers and anatomical scans.

Whole-brain anatomical data were collected for each participant with a resolution of 0.6 mm isotropic (MPRAGE sequences, FOV=229 × 229 mm^2^, matrix size=384 × 384, flip angle=5. T1-weighted: TR=3100 ms, TE=2.52 ms; proton-density-weighted: TR=1440 ms, TE=2.52 ms). For the functional runs, a 2D gradient-echo multi-band EPI sequence was used, with a resolution of 1.2 mm isotropic (multi-band acceleration factor=2 (Moeller et al., 2010), iPAT=3, FOV=172.8 × 172.8 mm^2^, matrix size=144 × 144, flip angle=75, number of slices=70, slice thickness=1.2 mm, no gap, ascending interleaved 2, TR=2000 ms, TE=21 ms, encoding direction Anterior to posterior, reference scan mode: GRE, MB LeakBlock kernel: off, fat suppression enabled). In each scanning session, a head scout was acquired for localization, then the B0 field map was acquired and loaded in the console. The interactive shimming was performed before acquiring the B1 field map. The system voltage was then computed according to the B1 map values (set to a maximum of 190 V across all sessions), to have a 90 degree flipping angle at the white matter beside the lateral ventricles, and the specific absorption rate (SAR) level for the longest functional run (432 volumes) was controlled at below 75%. The slices were tilted in an angle that covered most of the occipital lobe, parietal lobe and frontal lobe, while leaving out the anterior temporal lobe, part of the motor cortex, and the orbitofrontal cortex. This was to ensure that most of the important areas involved in body perception were covered, including EBA, fusiform gyrus, IPS, IPL, PMd, PMv. The amygdala was not consistently covered given the relatively limited coverage (covered in 6 out of 10 participants). Immediately before each functional run, a 5-volume run of the same setup but with posterior to anterior encoding direction was acquired (Invert RO/PE polarity: on), for post-hoc top-up EPI distortion correction (See the fMRI data preprocessing sub-section). We informed the participants the purposes of these distortion correction runs, and instructed them not to move between the distortion correction run and the actual functional run.

### Stimuli

The stimuli were gray-scale whole body images developed and validated in our lab (Stienen and de Gelder, 2011). They consisted of 8 actors, each posing 10 actions with or without emotional content. The first 5 categories were neutral actions: combing hair (CH), drinking water (DW), opening door (OD), talking on the phone (PH), putting on trousers (TR); the 6^th^ was neutral standing still (NE), and the last 4 were emotional expressions: fear (FE), anger (AN), happy (HA), sad (SA). The 80 postures were split into 2 balanced sets by randomly selecting images from 4 actors for each category, resulting in 2 sets of 40 stimuli (4 images per category, 5 images per actor), which ensures that in each stimulus category the participants perceive as much variability in the posture and the identity as possible. Each participant saw one of the sets.

The body stimuli were embedded in a gray background (RGB value = 128, 128, 128), with all internal facial information removed. They were sized to 400 × 600 pixels, and presented centrally on the screen (RGB value = 128, 128, 128). The whole-body shapes in the images overall spanned 309 × 492 pixels on the screen (visual angles=2.60 × 4.26 degrees).

### fMRI experiment design

The study used a slow event-related design. Stimuli were presented with Matlab (Version R2012a, the MathWorks, Natick, USA) and Psychtoolbox 3.0.11 (Pelli, 1997). A white fixation cross was present in the center of the screen throughout the experiment. The participants were asked to always fixate on the fixation cross and take in the body posture as a whole. The experiment consisted of 6 runs (14 min 18 s each, 429 volumes). Each run started with a fixation period of 8 seconds, and then the whole set of 40 stimuli was presented to the participant twice within each run. Each stimulus was presented for 500 ms, followed by an inter-stimulus interval of either 7.5, 9.5 or 11.5 s. The stimuli and the ISI were presented in a pseudorandomized order. In addition, 4 catch trials were included in each run. Within each catch trial, a body posture was randomly drawn from the stimulus set, while the fixation cross changed to either red or blue (RGB color red= 195, 32, 30; blue= 10, 109, 195) during the stimulus presentation period. Participants were asked to indicate the color by pressing the corresponding button of the button box as soon as they saw the fixation change color. Two seconds were added to the inter-stimulus interval after each catch trial. Excluding the catch trials, each single stimulus image was presented to the participant for 12 times throughout the main experiment. 9 participants completed all 6 functional runs; 1 participant completed 5 runs.

### Functional localizers

A separate scanning session was devoted to acquiring functional localizer data and the structural images. Stimuli were presented under passive viewing condition using Presentation software (Version 16.0, Neurobehavioral Systems, Inc., Berkeley, USA). In the static localizer run (14 min 24 s, 432 volumes), after a 12 s fixation period, gray-scale stimuli of faces, houses, bodies, tools and words were presented in blocks of 12 s (12 stimuli per block, 800 ms stimuli presentation, 200 ms inter-stimulus interval), followed by resting periods of 12 s with the fixation cross on a blank screen (RGB value=157,157,157). Each category block was presented 7 times, with a pseudorandomized presentation order for both the stimuli and the blocks. Facial stimuli were front-view neutral faces from the Karolinska Directed Emotional Faces (Lundqvist et al., 1998) (24 identities, 12 males). The part below the neck (clothes, hair etc.) was removed from the face images. Body stimuli were neutral still front-view bodies (de Gelder and Van den Stock, 2011) (20 identities, 10 males) from a different set than the one used in the main experiment, with the facial information removed. House and tool images were obtained from the internet. The house images consisted of 19 facades of houses with 2-to-3-storey height; the tool images consisted of 18 hand-held tools; words images consisted of high-frequency English words of 4-6 letters in Arial font. All the images were imbedded within a gray background (RGB value=157,157,157), spanning a visual angle of 1.99 degrees (230 pixels).

The dynamic localizer run (5 min 36 s, 168 volumes) consisted of 1-s video clips of either facial or bodily expressions, including neutral (coughing or clearing throat), angry, fear, happy (Kret et al., 2011). The actors performed the actions against a green background, either wearing black clothes for bodily expressions, or green clothes for facial expressions. Two exemplars were selected for each expression category (in total 8 for faces and 8 for bodies). The actors in the selected exemplars were all males. For the facial stimuli, two of the neutral expressions were performed by the same identity. The facial and bodily expression clips were presented separately in blocks of 8 s, with the stimuli order pseudorandomized. The face and body blocks were presented 10 times each, separated by an inter-block interval of 8 s, where a black screen and a white fixation cross was presented.

### Behavioral ratings

Immediately following the scanning session, participants completed a behavioral task outside the scanner. Each of the 40 stimuli participant saw in the scanner was presented once using Psychopy (v1.83.04)(Peirce, 2007) on an LCD monitor (Acer VG248, resolution = 1920×1080, refresh rate = 60 Hz, whole-body size in the stimuli spanning the visual angle of 13.42 × 8.43 degrees). For each stimulus image, 6 questions were answered either with a 7-point scale (for implied motion and valence respectively), or with an open answer with free typing (for the action the actor performed, and the emotion). See **Table S1** for details of the questions. We also recorded whether the participants changed their perception during the scan (questions 2 and 3). Four participants changed their perceptions (S4 and S9 for 2 stimuli, S6 for 7 stimuli, S10 for 12 stimuli). We used only the initial perception in the scanner (answer for question 1) for subsequent analyses. The same stimulus image stayed on the screen for all 6 questions. Participants answered the questions at their own paces (mean time spent=21.47 min, SD=6.98 min, range: 12.61 to 32.89 min).

### Data analysis

#### Representational similarity analysis (RSA) for behavioral data

For ease of use, we mapped the subjective reports with Deconf word embeddings (Pilehvar and Collier, 2016), which linked the Word2vec embeddings and the WordNet database. Word2vec embeddings were trained on a very large corpus of text, able to capture various linguistic relationships between words (Mikolov et al., 2013). After training, words with similar semantic meanings were found to be situated closer to each other in the embedding vector space (cosine distance); although for word2vec, different meanings of the same word were not disambiguated. On the other hand, WordNet (Miller, 1995) is a curated lexicon database, where different meanings of each single word were separated, and synonyms were grouped together; but it does not provide a quantitative mapping for word similarities. Deconf embeddings mapped the individual word meanings (senses) in WordNet into the 300-dimensional word2vec vector space, offering us both a common high-dimensional semantic space trained by a large corpus, and the precise separation of different meanings.

57.8% of the behavioral free reports collected were single words (action: 110/400; emotion: 353/400). For phrases and sentences, we omitted the pronouns (e.g. he is, she is, his, her), as most of the times participants did not consistently type in the pronouns. The rest of the words/phrases (nouns, verbs, adjectives, adverbs) that had a corresponding entry in WordNet 3.1 were lemmatized (e.g. stretching→stretch). For words/phrases not found in WordNet 3.1, “how” was omitted; “something” was substituted by “thing”; “himself/herself/oneself” were substituted by “self”. The adverb “just” associated with verb phrases was omitted (e.g. just watching→watch). When describing the perceived action, “not sure” was substituted by “unsure”; when denoting that the participant had no idea about what the person was doing, “no idea” and “not sure” were substituted by “not applicable”. For words with multiple meanings (“senses” the term used in WordNet), the corresponding sense was selected. When a noun denoting emotion (e.g. “happiness”) has both senses of <noun.feeling> and <noun.state>, the sense of <noun.feeling> was always selected, since this sense was the one mentioning “emotion”. There were very few cases that participants typed in “not”. For one case the participant denoting the emotion of the person in the stimuli “was not sure about something”, it was substituted with the word “unsure”. All the other cases were in responses for perceived actions, about participant’s own uncertain understanding for the stimuli. For these and similar cases we substituted the entry with “not applicable”. For all response entries and word lists, see https://osf.io/cuh9v/?view_only=efb12b7585ee4b6bbcfd7ca42c63b60d.

For each free report entry after lemmatization, the 300-dimension word vector with the corresponding sense number and sense key were selected from the Deconf pre-trained embeddings. When an entry has multiple words, the vectors were averaged for this entry. The RDM for each participant were then computed in cosine distance (reflecting angles between vectors), as this metric was routinely used for computing word-embedding distances in the literature. Averaging resulted in the same matrix as addition (the angles didn’t change).

The RDMs for behavioral ratings (implied motion, valence) were computed directly from the ratings, in Euclidean distance.

For predefined RDMs, each category was binary-coded in vectors, with numbers of elements corresponding to the total number of categories. E.g. for predefined actions, drinking water (2^nd^ category in 10) was coded as 0 1 0 0 0 0 0 0 0 0; for predefined non-emotion/emotion categories, emotional ones were coded as 0 1. This coding assumes that each category was orthogonal to the others. The resulting RDMs were computed in Euclidean distance.

For RDM comparisons throughout the study, the Spearman’s correlation was computed between 40×40 RDMs, with the 780 values below the diagonal of the RDMs as inputs. The resulting rho values were Fisher’s Z transformed, submitted to a one-sample t test against 0 (two-tailed) at the group level. The group-averaged Z values were back transformed to rho values (or 1 - rho distance) and reported. To assess individual variability, we computed the coefficient of variation (CV, sample standard deviation/sample mean, in %).

Since all 400 response entries across participants for the perceived action and emotion representations were in the common 300-dimensional word-vector space, we performed PCA on the 400 entries respectively for perceived action, and for perceived emotion (without centering, since the word vectors were all normalized). To see whether predefined categories were reflected in some dimensions of the subjective representations, the PCA scores for each principal component (PC) was computed into RDMs (Euclidean distance), and correlated with 400×400 model RDMs. The model RDMs were created in the same way as the 40×40 RDMs, with the only difference that the 400 entries were sorted by action categories across all participants. Model RDMs included predefined action, predefined emotion, implied motion, valence, individual subjects (entries of the same participant were coded as the same vector). The PCA scores of all response entries were plotted in the two PCs that showed the highest positive correlation to the model RDMs (**Fig. 1I, J**).

### fMRI data Preprocessing

The data were processed with BrainVoyager (version 20.0, 20.2, 21.45, Brain Innovation, Maastricht, the Netherlands), MATLAB (version R2016a), and NeuroElf 1.0 toolbox implemented in MATLAB (http://neuroelf.net/). Before any preprocessing, the functional data underwent in-plane EPI distortion with the COPE plugin (v1.0) in BrainVoyager (https://support.brainvoyager.com/brainvoyager/available-tools/86-available-plugins/62-epi-distortion-correction-cope-plugin). The voxel-wise displacement was estimated between the first volume of each functional run and the first volume of the preceding 5-volume correction run (in reversed phase encoding direction). The in-plane voxel-wise displacement map was then applied to all the volumes of the functional run. The first volume of each run was saved as a separate file, and applied with distortion correction. This volume then served as the basis for subsequent motion correction and across-run alignments. After distortion correction, the functional runs underwent slice scan time correction with the slice timetable information from the scanner (interpolation: sinc), within-run 3D rigid motion correction with each run’s first volume as references (interpolation: trilinear for motion estimation, sinc for applying transformations), and temporal high-pass filtering (GLM with Fourier basis set of 2 cycles, including linear trend). The order of distortion correction and motion correction was assessed on S06’s sixth (last) main-task run, which has the biggest within-run rotation in all acquired runs in the current experiment (about 3 degrees in the x axis). Between the two processing orders, the resulting 429^th^ volume showed negligible differences in the frontal lobe. Thus we believe the order of distortion correction and motion correction was not critical. For the anatomical data, the magnetic field inhomogeneity was corrected by dividing the T1 images with the PD images. The anatomical data was then spatially normalized into the Talairach space.

To ensure that all functional runs of each participant (6 runs of the main experiment, 2 functional localizer runs, from two scanning sessions, distortion-corrected) were aligned well with each other, we used a manual across-run alignment procedure, with careful visual inspections and multiple iterations/checks. The first volume of the main-task run was aligned to the anatomy, and then saved as an anatomical file in native space, keeping the T2* weighted contrast the same as the original functional run. This served as a “dummy” anatomical run for the subsequent alignment procedure. Then the first volumes of all the functional runs (including the first run itself) were aligned to this dummy anatomical run, and normalized into the Talairach space (with the position information and transformation matrices for the T1 weighted anatomical run). The first volumes of the 8 runs aligned in the Talairach space were then screen-captured and saved as different layers in Photoshop (version CS6, Adobe, United States), toggled on and off, and made into GIF animations to check alignment qualities across runs. Adjustments were made for imperfect aligned runs, followed by the same checking procedure, until there were not any big shifts/translations across runs at the whole-brain level. This procedure allowed us to visually spot tiny alignment imperfections across runs with the same T2* weighted modality. It is much easier than spotting misalignments across T2* weighted and T1 weighted image modalities in the conventional alignment procedure, where the two images look very different across modalities. The quality of the distortion correction was also checked when aligning the functional run 1 to the anatomy, where a good distortion correction resulted in a good shape correspondence between the T2* weighted and T1 weighted data. In some cases, when the participants moved their head between the distortion correction run and the functional task run, despite our repetitive instructions, the distortion correction quality was affected. In those cases, we strived for aligning the occipital and temporal lobes.

After alignment across runs, the functional data of the main experiment were then spatially smoothed with 6 mm FWHM for univariate analysis (comparable to the RSA searchlight radius), 3 mm FWHM for task residual functional connectivity analysis. Due to the individual anatomical/functional variability, 3 mm was chosen for the functional connectivity analysis, as a compromise between the 6 mm smoothing and no smoothing, with the former rendering too extensive connectivity patterns almost covering the whole brain (data not shown), and the latter rendering too much false negatives (shown in the functional localizer analysis). The unsmoothed data were used for the (multivariate) representational similarity analysis, and univariate analysis of 10 categories in RSA searchlight clusters. The static functional localizer data was smoothed at 6 mm FWHM for group random-effect GLM, 3 mm for individual participants, and not smoothed for univariate analysis in RSA searchlight clusters.

All results in this study were computed in the volume space. For visualizing group-level results, cortex-based alignments (CBA) were performed to alleviate the high inter-individual anatomical variability. For each participant, the gray-white-matter boundary of the anatomical images in Talairach space underwent automatic segmentation, and careful manual correction slice-by-slice. Then a mesh with a high number of vertices was created for the white-gray-matter boundary of each hemisphere (number of vertices ranging from 370k to 434k), inflated into a sphere, corrected for distortions across vertices, and mapped to a high-resolution standard sphere with down sampling (number of vertices=163842). The curvature patterns of the original mesh were also smoothed and mapped to the standard sphere. The group-level cortex-based alignment was separately performed for the left and right hemispheres. For each hemisphere, the curvature patterns of all 10 participants’ meshes were aligned into a dynamic group average. After aligning, the average left and right hemispheres of the whole group were created. This resulting group-average brain surface retained many anatomical landmarks, and was more detailed than the ones usually created with 3T data. However, note that it was a bit smaller than the actual brain, and was only meant to provide visual landmarks to localize the clusters.

### Univariate analysis

To compare the univariate activation to the literature, three different general linear models (GLMs, serial correlation correction: AR(2), data %-normalized before performing the GLM) were applied to the datasets smoothed 6 mm FWHM, with the following predictors: 1) 10 action categories; 2) the behavioral ratings of implied motion; 3) the behavioral ratings of valence. For 2) and 3), the presentation of all stimuli were defined as a single main predictor, the ratings for 40 stimuli were z-scored within each participant, and served as a parametric weight factor to each corresponding stimulus. The parametric modulation effects were subsequently computed for 2) and 3). For all the GLM models above, the time courses were %-transformed, the main predictors were convolved with a two-gamma hemodynamic response function, and the 6 parameters of participant’s head motion were z-scored and entered as confounding factors. The catch trials were marked as a separate condition, and their parametric estimates (beta values) were not used in subsequent analyses.

The group random-effect GLM analysis was performed for each predictor set. For the 10-category predictor set, apart from the contrasts between action categories and versus baseline, a whole-brain ANOVA was performed, with a 10-level factor “action categories”. Cluster size thresholds for all group-level maps in this study were estimated using Monte-Carlo simulation (alpha level=0.05, numbers of iterations=5000, initial p<0.005 for univariate and RSA searchlight analyses; initial p<0.001 for task-residual functional connectivity), with the BrainVoyager plugin Cluster-Level Statistical Threshold Estimator (https://support.brainvoyager.com/brainvoyager/functional-analysis-statistics/46-tresholding-multiple-comparisons-problem/226-plugin-help-cluster-thresholding), masked with the common functional data coverage across 10 participants. This mask was created from the averaged functional images across participants, covering 830178 functional voxels.

The GLM of the static functional localizer (without smoothing, and smoothed 3 mm FWHM) was performed for each individual participant. Since we do not assume that the bodies should only be processed in category-specific areas, the static functional localizer data were used as an indicator for inter-individual variability, and an indicator for the neural processes in some RSA searchlight clusters.

### Representational similarity analysis for fMRI data

For RSA analyses, the GLM model with 40 stimuli was fitted to each individual participants’ unsmoothed data, resulting in 40 t-maps, one per stimulus. The neural RDMs across the 40 stimuli were computed with Pearson’s correlation distance. Searchlight spheres were constructed for each voxel, with a radius of 5 voxels (515 voxels within the sphere, 889.92 mm^3^), comparable to the value for univariate smoothing (6 mm). Spheres containing more than 172 non-zero voxels were included in the analysis). The Spearman correlation values between the neural RDM and the model RDM was Fisher-Z transformed, and wrote back to the sphere center voxel. The group-level significance of the correlation was evaluated by a one-sample t-test against zero (2-tailed).

For the clusters found by the searchlight RSA and subsequent whole-brain analyses, the univariate percent signal changes for each of the 10 action categories were extracted (without smoothing), and were compared to the baseline (one-sample t test against 0) in SPSS.

### RSA regression with low/mid-level features

For each of the stimuli image, we extracted the body joint locations by the OpenPose library (Cao et al., 2019) (https://github.com/CMU-Perceptual-Computing-Lab/openpose). The output was the x, y values and a confidence score for the estimation, for 18 body joints (See **Figure 4A**). The joint locations were imported in Adobe Illustrator CS6, overlaid on the stimulus image, and manually adjusted with visual estimation. The correspondence of the joints and their location are as follows: joint 1 (head)-nose tip; joint 2 (neck)-the jugular notch of the sternum; joint 3 & 6 (shoulders)-humerus; joint 4 & 7 (elbows)-the connecting point of the upper/lower arms; joint 5 & 8 (hands)-the wrist end of the radial bone; joint 9 & 12 (waist)-the widest points of the femur; joint 10 & 13 (knees)-on the patella; joint 11 & 14 (feet)-the ankle end of the tibia; joint 15 & 16 (eyes); joint 17 & 18 (ears). Since the eyes were not present on the stimuli images (facial features blurred), and the ears were often occluded by the hair or by the head, these joints were visually estimated with linear perspective in mind.

We constructed low-level visual feature RDMs, for raw pixel values (as vectors), and raw coordinates of the 14 joints (excluding joints for ears and eyes), although the resulting RDMs were extremely similar to the ones including these 4 joints (RDM similarity rho=0.976, 0.98 for stimuli set A and B).

The mid-level visual feature RDMs included head, shoulder, waist orientations, as well as hands/feet-to-head distances. The head orientation was individually computed by the distance between ears, normalized (divided) by the ear distance of the standing still condition, capped at 1 (1: facing the viewer; smaller than 1: facing left- or right-ward). The shortest hand-to-head distance may reflect whether the hand enters the head’s peripersonal space of the actor; the average feet-to-head distance may reflect the relaxation of the torso and legs of that actor.

The low- and mid-level visual features were chosen for their theoretical and behavioral relevance, and their ease for computing/handling. These were only a small subset in an infinite model space.

For RSA regression in each of the 10-action-category areas (resulted from whole-brain searchlight analysis of the 10-action-category model RDM), the low/mid-level visual feature RDMs and the higher-level stimuli attribute RDMs (implied motion, valence, actor identity) were squared for each element (to satisfy the linearity for squared Euclidean distance values), z-scored across elements, then put together in the same linear regression model as predictors. Each element of the neural RDM was also correspondingly squared. A set of beta values for these predictors were obtained for each participant, and their group-level significance was examined by one-sample t-test against zero (2-tailed), and FDR-corrected at q<0.05 (Storey, 2002) across all predictors for each searchlight cluster.

### Task residual functional connectivity analysis

We smoothed the task data at 3 mm FWHM, regressed out the task-related activity by deconvolution analysis (5 stick predictors per stimulus, covering the evolvement of the BOLD shape per trial). For each seed region, the residual time course was extracted and averaged across voxels, and correlated with the residual time courses of all voxels in each run, resulting in one R map per run. The connectivity pattern across the runs were stable for all participants. The R maps were Fisher’s Z-transformed, averaged across runs per participant, and put in a one-sample t-test against 0 (2 tailed). The resulting group-level maps were thresholded at p<0.001 for cluster size correction.

For the 10-action-category areas, the inter-area functional connectivity was also examined, by correlating the seed ROI time courses per run, then again with the same group-analysis procedure (Z-transform, averaging, t test), with FDR correction.

### Hierarchical clustering of RDMs across functional connectivity clusters

We computed neural RDMs for all group-level clusters obtained from the task-residual functional connectivity analysis (51 ROIs for the perceived action FC network, 76 ROIs for the perceived emotion FC network), and computed second-level RDMs (Spearman correlation distance, **Figure 6F**) from them within each participant. For the two FC networks, the second-level RDM were averaged (with Z-transform) across participants (**Figure 6A, B**, right panels), and served as inputs for hierarchical clustering (MATLAB function: linkage; method: average. Plot function: dendrogram; clustering threshold for the most similar areas to the seed region: 0.8). We also examined the univariate activation for 10 action categories, and plotted histograms.

## Acknowledgments

This study was supported by the European Research Council, under the European Union’s Seventh Frame-Work Programme (FP7/2007–2013)/ERC (grant numbers 295673 to BdG and 269853 to RG), by the ERC-Synergy program grant RELEVANCE (Grant agreement 856495 to BdG), and by the FPN-MBIC funding of Maastricht University to MZ and BdG. MZ was supported by Fondation Bettencourt Schueller. We thank Federico de Martino for setting up the 7T scanning sequences and Giancarlo Valente for providing a part of the searchlight code. We thank Maarten Vaessen for commenting on a previous version of the manuscript.

## Author Contributions

**Conceptualization**: M.Z. and B.dG.; **Methodology, Software, Validation, Formal Analysis, Investigation, Data Curation, Writing – Original Draft, Visualization**: M.Z.; **Resources**: R.G.; **Writing – Review & Editing**: M.Z., R.G. and B.dG.; **Project Administration, Funding Acquisition**: M.Z. and B.dG.; **Supervision**: B.dG.

## Data and code availability

The data and codes of this study are available at https://osf.io/cuh9v/?view_only=efb12b7585ee4b6bbcfd7ca42c63b60d, including: stimuli images with body joint estimations, subjective reports and corresponding word embeddings, whole-brain statistical result maps, ROI raw data for the RSA analysis, structural images and white-gray-matter-boundary segmentations.

