## Supplementary materials for "Subjective understanding of actions and emotions requires interaction of the semantic and action observation networks"

### Supplementary Information

#### Supplementary figures

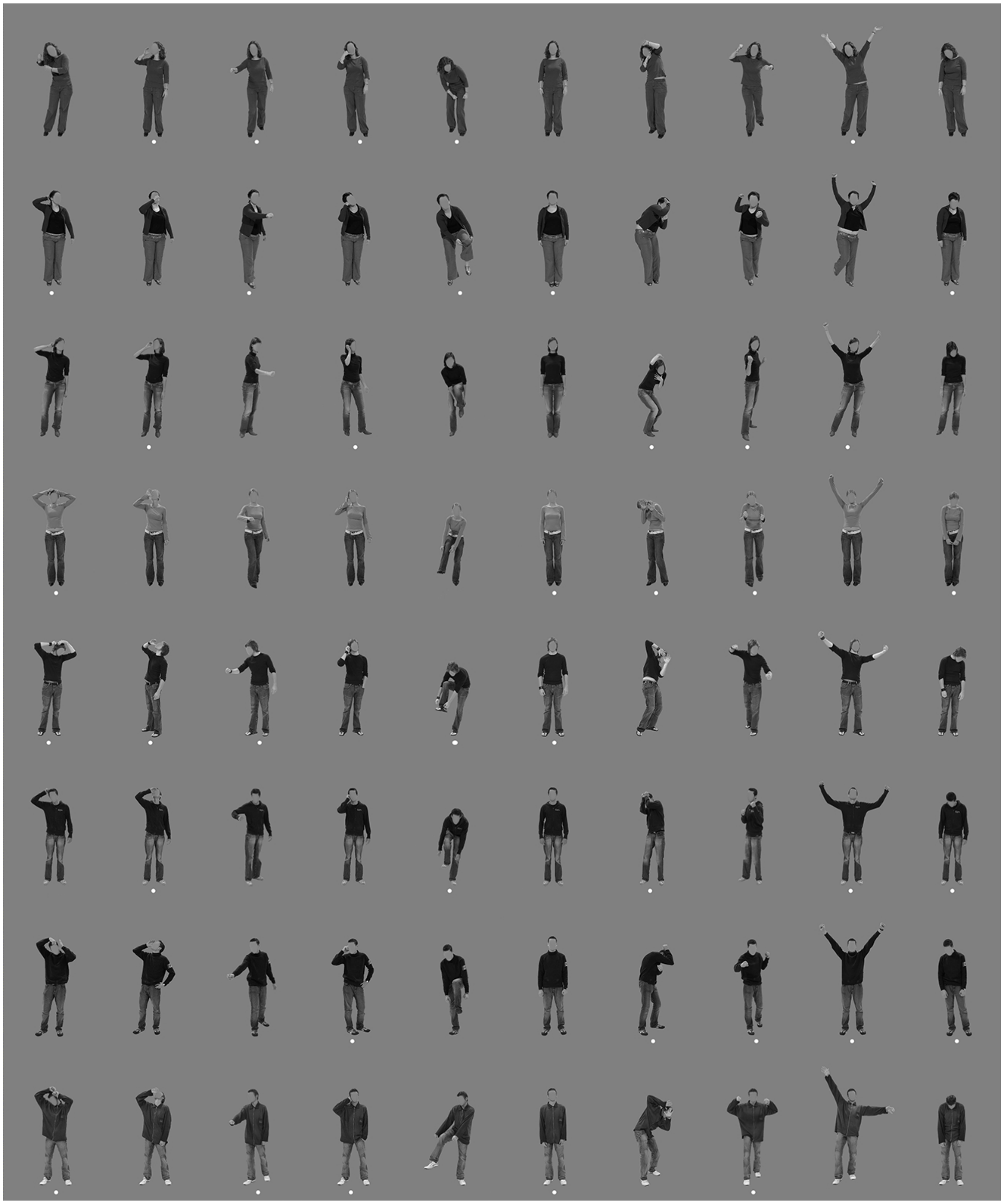

**Figure S1.** All of the stimuli images, which comprised of 10 action categories, performed by 8 actors. The stimuli were separated into two balanced sets. The ones marked by dots were in set A, presented to S01 to S05. The other half of the stimuli were in set B, presented to S06 to S10.

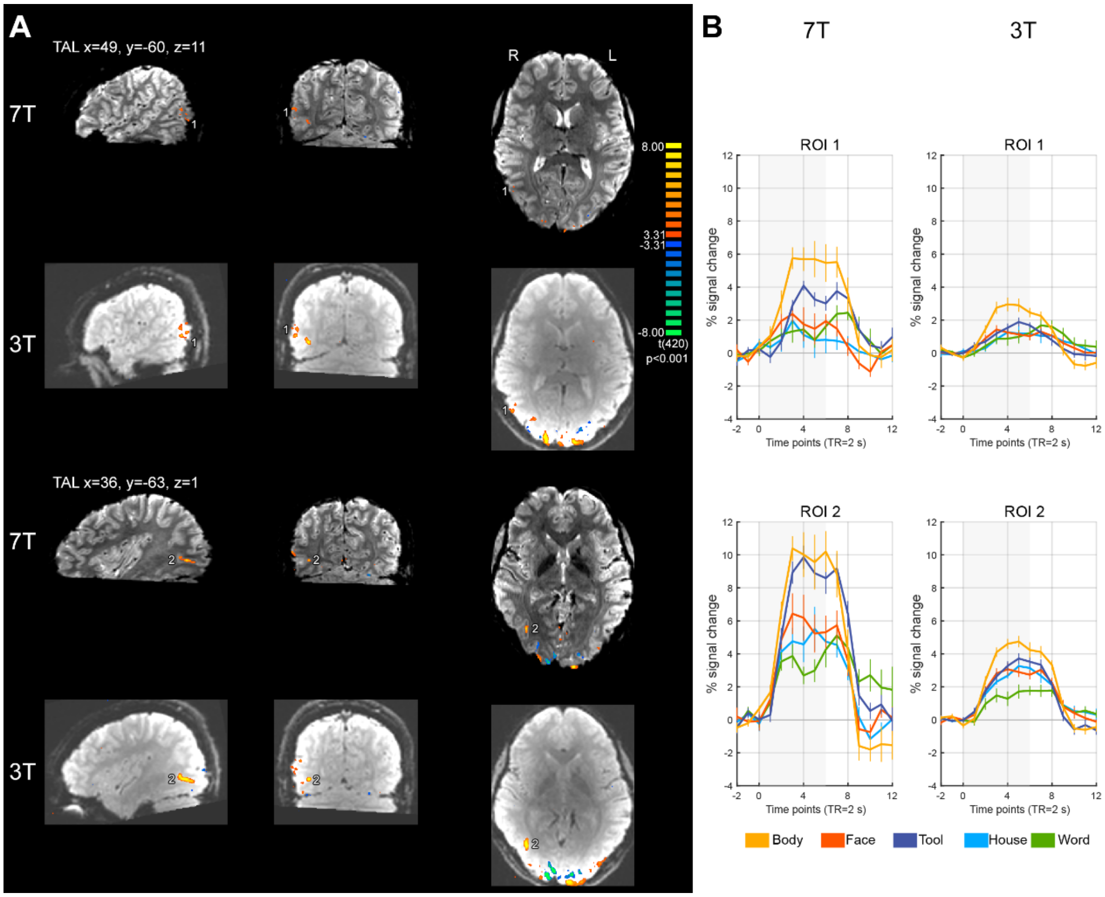

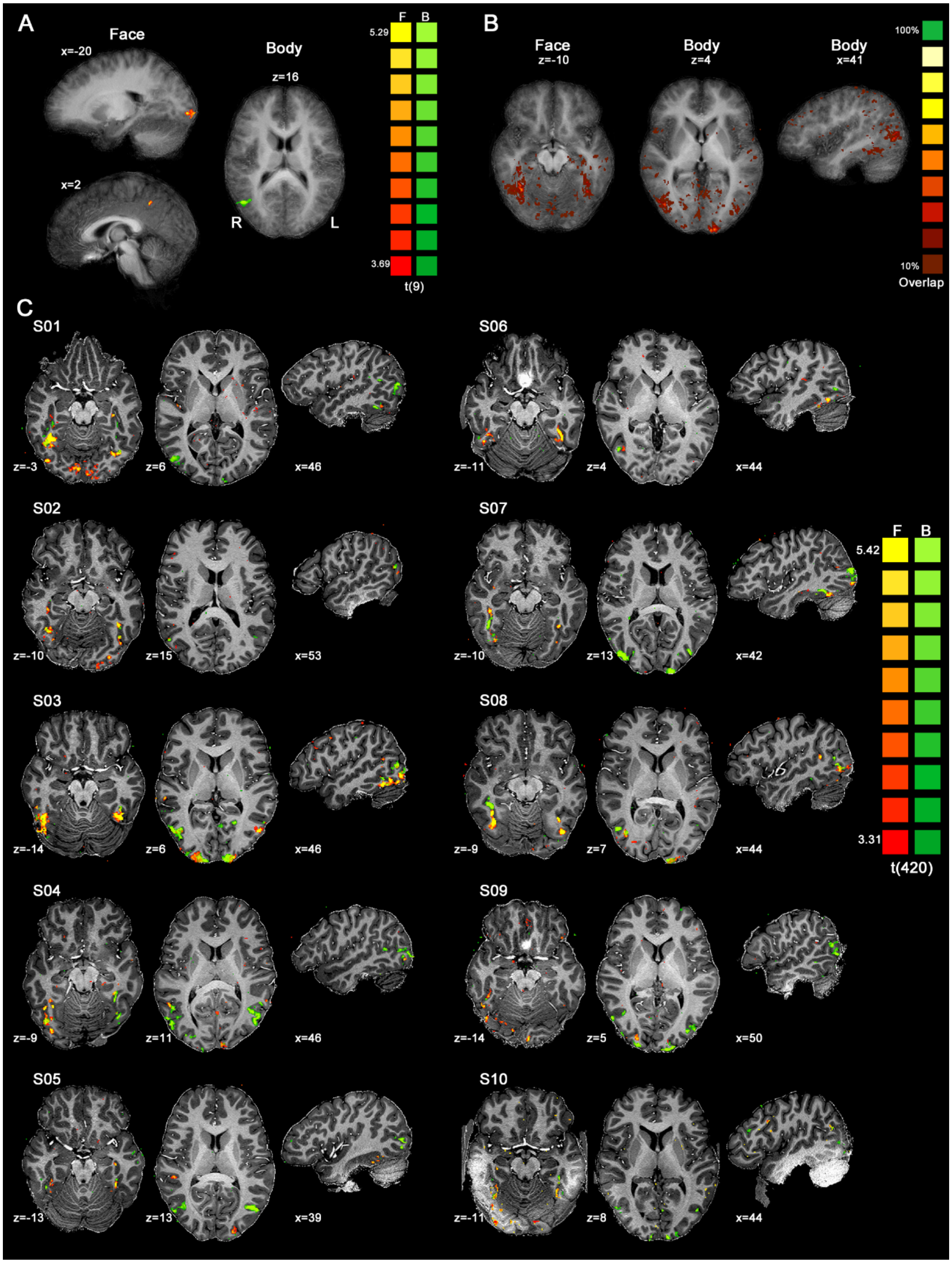

**Figure S3.** The inter-individual variability, shown in the functional localizer data. **A.** Group-level activation maps for faces>other 4 object categories, and bodies>other 4 object categories (data smoothed 6 mm FWHM, Monte-Carlo simulation, initial p=0.005, alpha=0.05, n simulations=5000, face cluster threshold=66, body cluster threshold=70). Color bar p range: 0.005 to 0.0005. **B.**

Overlap of individual face- and body-specific clusters across participants was very low, both ranging from 10% to 50% (reaching 50% in very few voxels). **C.** Robust face- and body-specific clusters in each individual participant (data smoothed 3 mm FWHM, p=0.001, uncorrected). Color bar p range: 0.001 to $1\times10^{-7}$. The anatomical data quality at occipito-temporal lobes for S10 (without dielectric pads) was worse than the other participants (with dielectric pads).

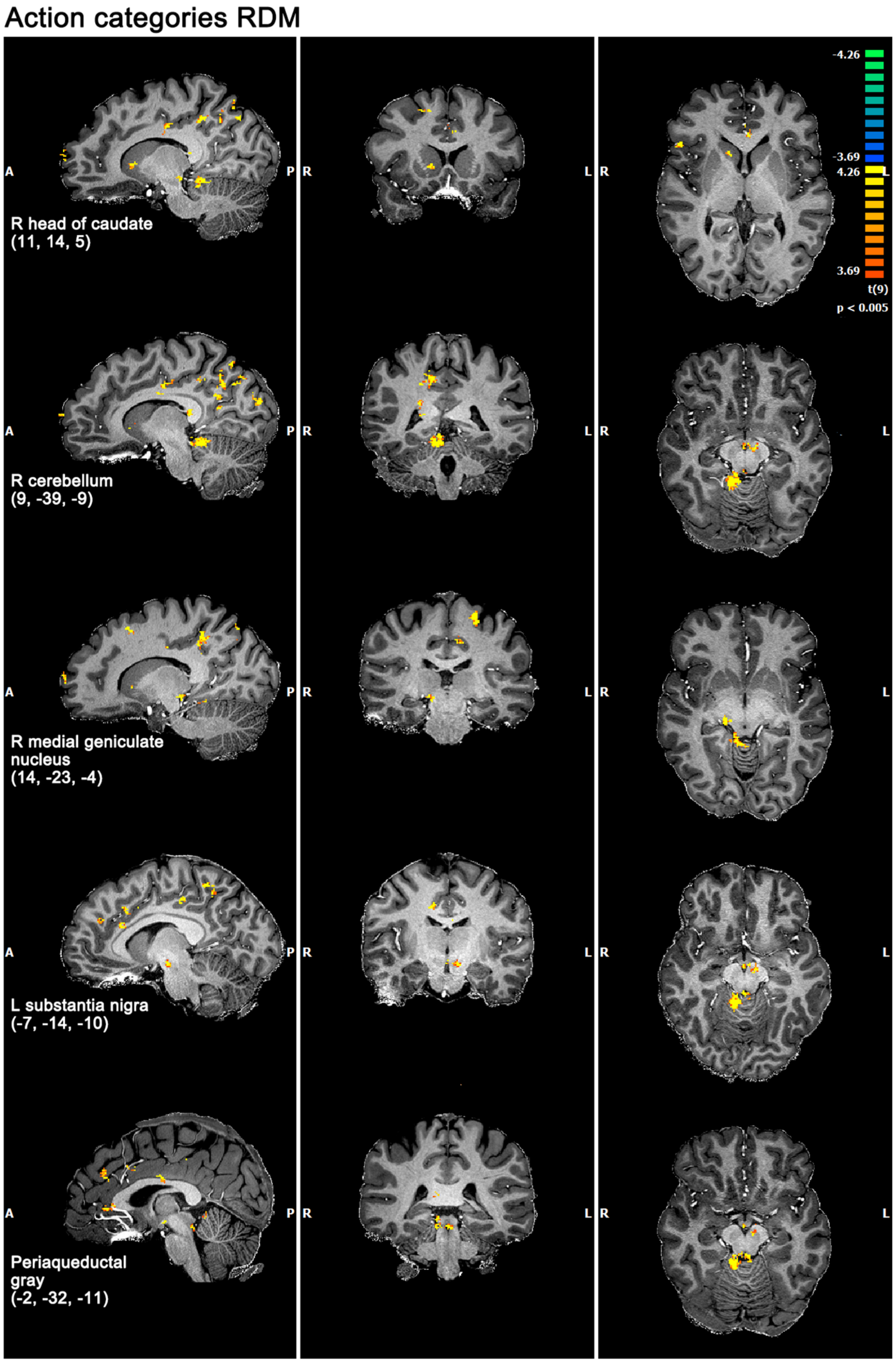

**Figure S4.** Subcortical and cerebellar clusters showing significant correlations to the 10-action-category RDM (alpha=0.05, initial p=0.005, cluster size corrected), plotted on the T1 image of a representative participant in volume space. The Talairach coordinates of each view are shown in brackets. Color bar p range: 0.005 to 0.0021.

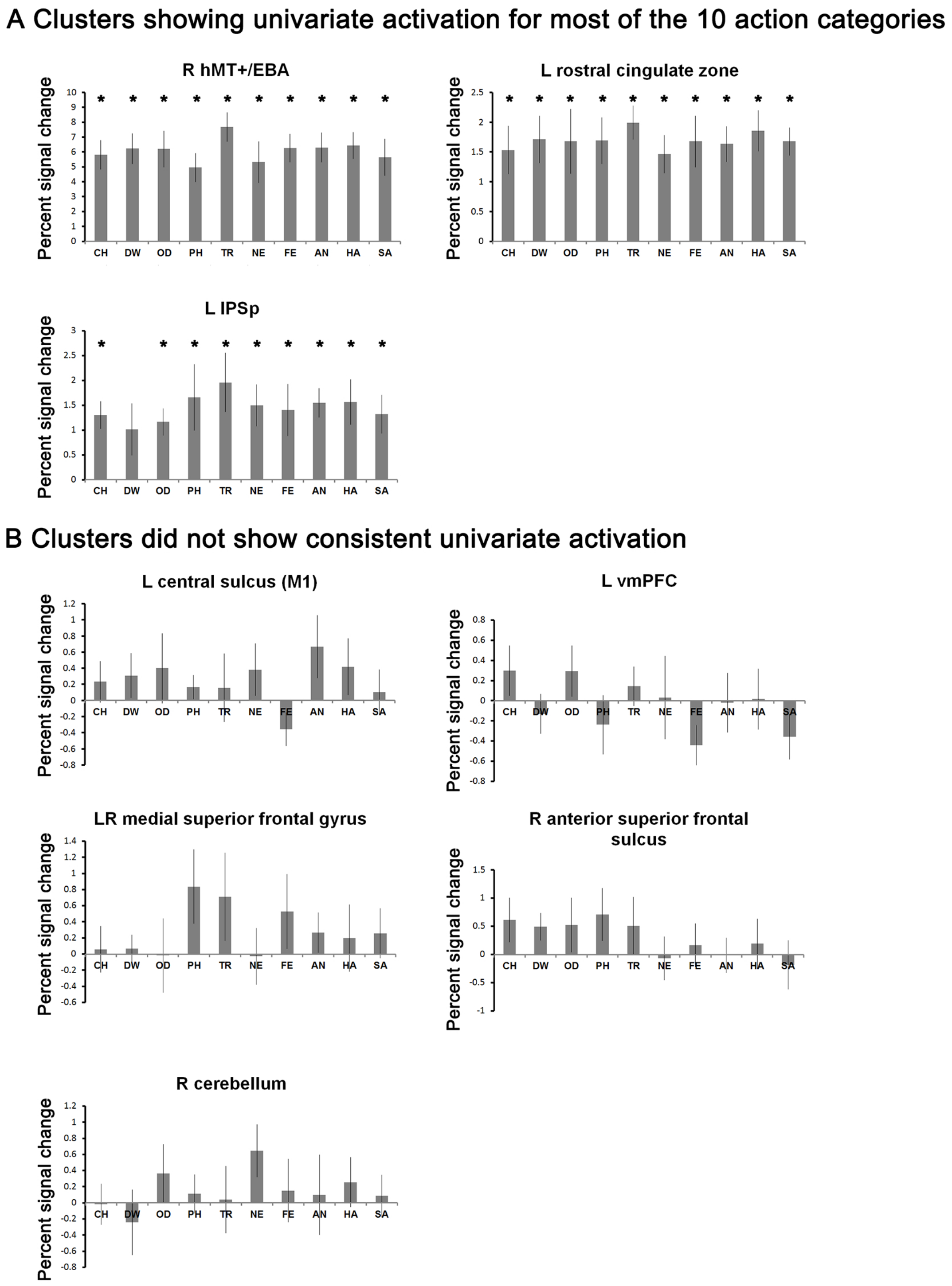

**Figure S5.** The univariate activation of the 10 individual action categories in the 10-action-category RDM clusters, comparing to the baseline (fixation cross). **A.** Clusters showing consistent univariate activations at the group level. * denotes the percent signal change significantly bigger than 0. **B.** Clusters did not show consistent univariate activations. Note that the percent signal changes in the 7T functional data were much higher than the ones found in 3T studies with an event-related design, see the right hMT+/EBA cluster in **A** for an example. Error bars denote SEM.

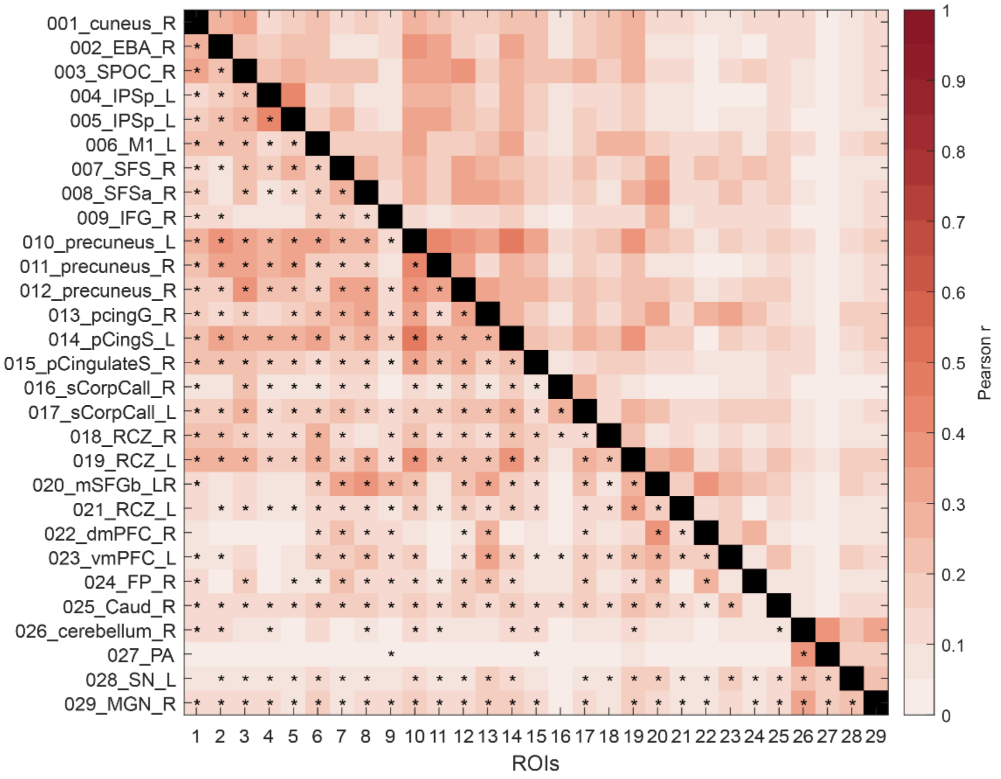

**Figure S6**. Functional connectivity between the 10-action-category areas. FDR q<0.05 for the connectivity pairs below the diagonal. Significant connectivity pairs were marked with an “*”.

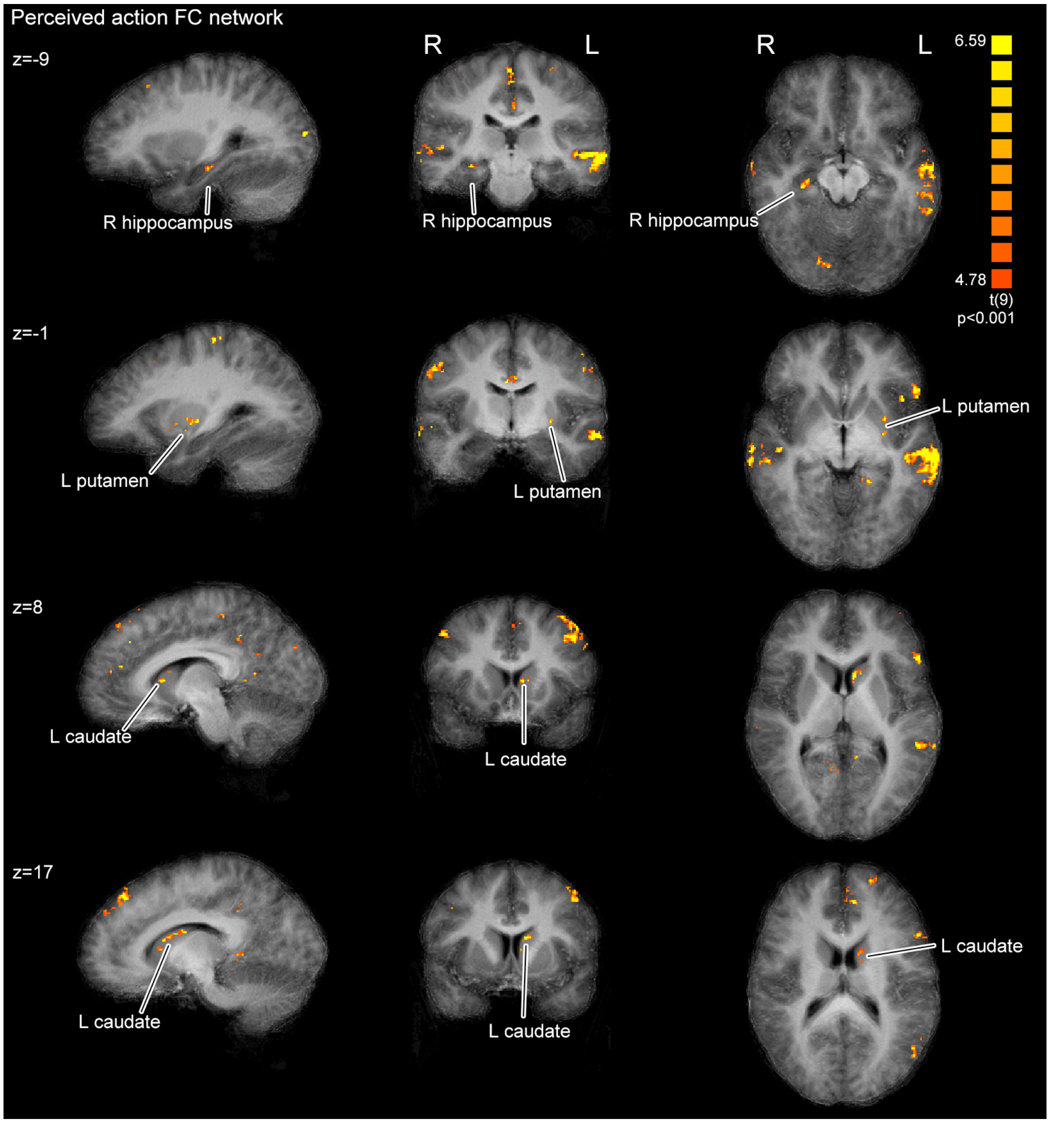

**Figure S7.** Subcortical, cerebellar and hippocampal regions observed in the perceived action FC network, plotted on the T1-weighted anatomical images averaged across 10 participants (Fisher's Z-transformed Pearson correlation maps, one-sample t-test against 0, cluster size thresholded at alpha=0.05, initial p=0.001, Monte-Carlo simulation n=5000. Cluster size threshold=32 functional voxels). Color bar p range: 0.001 to 0.0001.

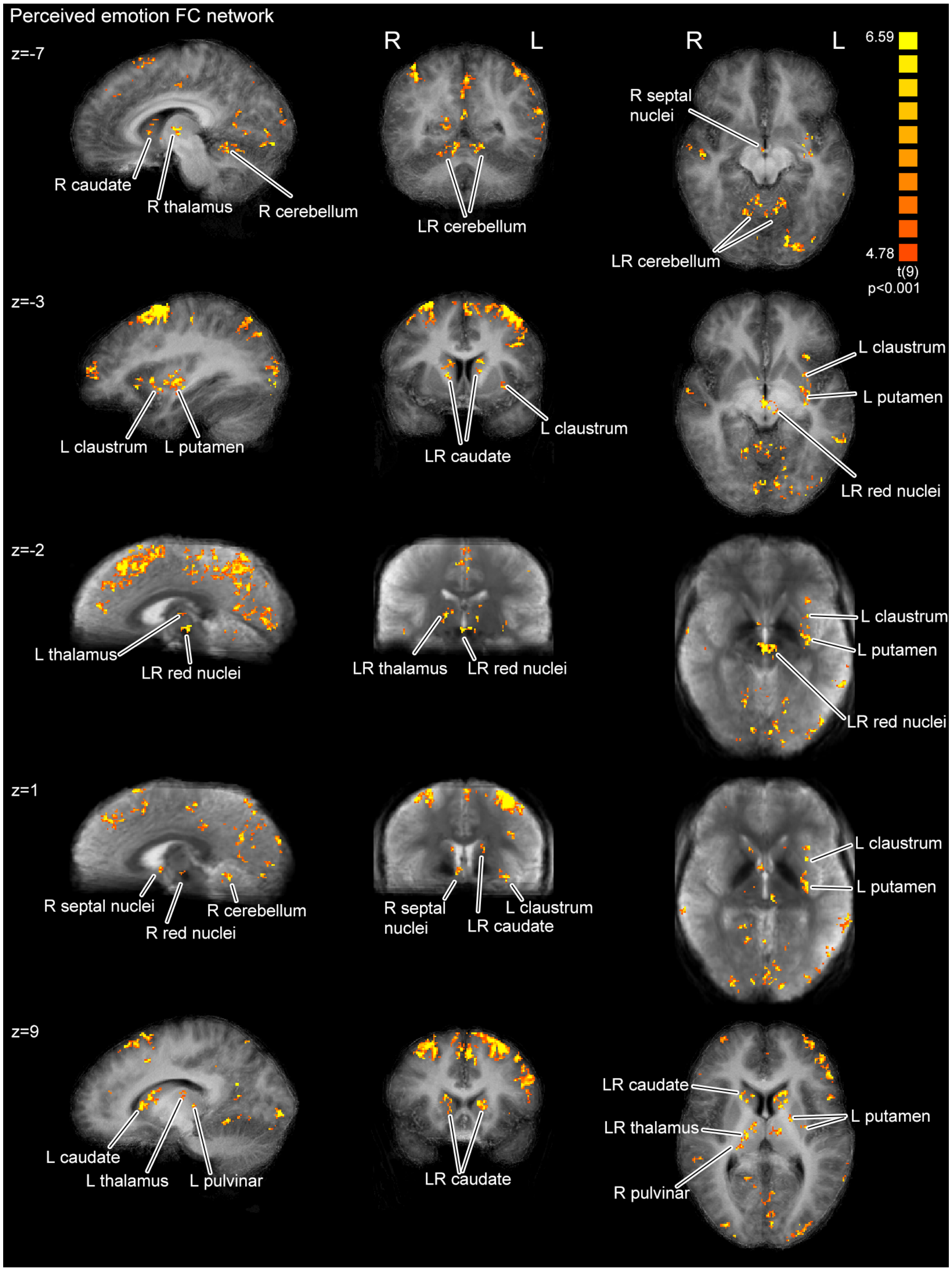

**Figure S8.** Subcortical, cerebellar and hippocampal regions observed in the perceived emotion FC network, plotted on the T1-weighted anatomical images averaged across 10 participants (Fisher's Z-transformed Pearson correlation maps, one-sample t-test against 0, cluster size thresholded at alpha=0.05, initial p=0.001, Monte-Carlo simulation n=5000. Cluster size threshold=40 functional voxels). Color bar p range: 0.001 to 0.0001. The bilateral red nuclei and the globulus palidus could be visually distinguished by the T2*-weighted contrast, shown as dark areas in images of z=-2 and z=1.

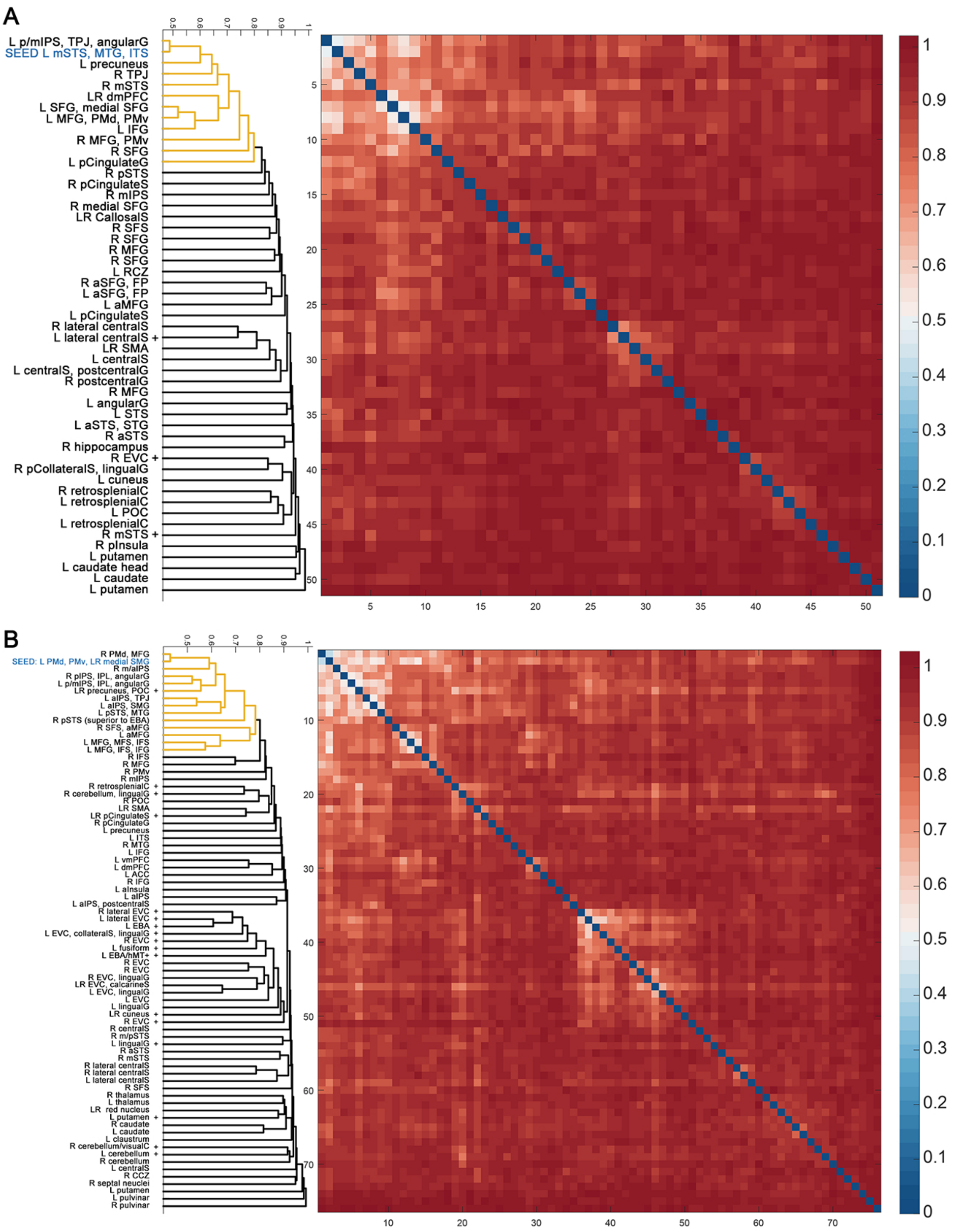

**Figure S9.** Corresponding to Figure 6, for the clusters found in the group-level task-residual functional connectivity analysis, the dendrograms and group-averaged second-level RDMs for the multivariate patterns in these clusters were plotted together here.

#### Supplementary tables

**Table S1.** Questions and types of answers in the behavioral task after the scan. *Participants were instructed to type "na" when not applicable. See all subjective reports in the data available online (https://osf.io/cuh9v/?view_only=efb12b7585ee4b6bbcfd7ca42c63b60d).

| Questions | Types of the required answer | Example answers to stimulus F3HA (Figure S1, row 3, column 9) |
|---|---|---|
| 1) What did you think this person was doing, when you saw this picture in the scanner? | Free report, typing | S03: cheering<br>S05: yawning |
| 2) Did your opinion on the activity he/she was doing change during the scan? | Selection: changed/remained the same | S03: Remained the same<br>S05: Remained the same |
| 3) If changed, what did you think he/she was doing initially, and later? | Free report, typing | S03: na*<br>S05: na |
| 4) Please rate the amount of movements present in the image, with a scale of 1 to 7. | Rating | S03: 1<br>S05: 3 |
| 5) Please rate the valence of the body picture, with a scale of 1 to 7. | Rating | S03: 7<br>S05: 6 |
| 6) If you think the person was expressing emotions, which emotion do you think it is? | Free report, typing | S03: being happy<br>S05: relaxation |

**Table S2.** Comparison of emotion reporting accuracy (%) in the current study, to two validation studies of facial emotional databases.

| Study | N participants | Accuracy measure | Neutral | Fearful | Angry | Happy | Sad |
|---|---|---|---|---|---|---|---|
| Current study | 10 | Percentage of free-report responses corresponding to the intended emotion | 75 | 87.5 | 87.5 | 82.5 | 52.5 |
| Langner et al. (2010), Radboud Faces Database | 276 | Forced choices, percentage of chosen emotions per intended emotional expression | 84 | 83 | 85 | 98 | 80 |
| Goeleven et al. (2008), Karolinska Directed Emotional Faces | 272 | Forced choices, percentage of participant who correctly identified the emotion (biased hit rate percentage) | 62.64 | 43.03 | 78.81 | 92.65 | 76.70 |

**Table S3.** Clusters found in the searchlight RSA, for the 10-action-category model, the non-emotion/emotion model, and for attribute models (alpha=0.05, initial p=0.005, cluster size corrected). The R values were back transformed from Fisher's Z values. Subcortical and cerebellar clusters were listed after the cortical ones. * denotes clusters that survived thresholding at the initial p=.001.

| Reference RDM | Location | Hemi sphere | initial p=.001 | cluster size (mm3) | average R | SD R | average t | average p | TAL x | TAL y | TAL z |
|---|---|---|---|---|---|---|---|---|---|---|---|
| **Action categories** | cuneus | R | * | 224.6 | 0.026 | 0.011 | 4.747 | 0.002 | 7 | -82 | 24 |
| | hMT+/EBA | R | * | 140.0 | 0.028 | 0.013 | 4.671 | 0.002 | 49 | -55 | 11 |
| | SPOC | R | | 172.8 | 0.018 | 0.007 | 4.651 | 0.002 | 9 | -69 | 40 |
| | IPSp | L | | 124.4 | 0.022 | 0.008 | 4.538 | 0.002 | -25 | -50 | 37 |
| | IPSp | L | | 76.0 | 0.019 | 0.007 | 4.537 | 0.002 | -26 | -60 | 41 |
| | central sulcus (M1) | L | * | 286.8 | 0.029 | 0.009 | 4.720 | 0.002 | -22 | -25 | 58 |
| | superior frontal sulcus | R | | 160.7 | 0.023 | 0.011 | 4.761 | 0.002 | 18 | 16 | 47 |
| | superior frontal sulcus, anterior cluster | R | | 114.0 | 0.028 | 0.016 | 4.309 | 0.003 | 21 | 32 | 34 |
| | inferior frontal gyrus | R | | 91.6 | 0.023 | 0.009 | 4.879 | 0.002 | 50 | 22 | 5 |
| | precuneus | L | * | 409.5 | 0.021 | 0.006 | 4.603 | 0.002 | -11 | -48 | 49 |
| | precuneus | R | | 74.3 | 0.020 | 0.010 | 4.307 | 0.003 | 11 | -64 | 51 |
| | precuneus | R | | 179.7 | 0.025 | 0.011 | 4.462 | 0.002 | 9 | -55 | 38 |
| | posterior cingulate gyrus | R | * | 482.1 | 0.025 | 0.009 | 4.502 | 0.002 | 5 | -52 | 20 |
| | posterior cingulate sulcus | L | | 146.9 | 0.020 | 0.007 | 4.671 | 0.002 | -8 | -25 | 41 |
| | posterior cingulate sulcus/caudal cingulate zone | R | * | 343.9 | 0.023 | 0.008 | 4.603 | 0.002 | 14 | -40 | 40 |
| | sulcus of corpus callosum | R | | 108.9 | 0.020 | 0.006 | 4.546 | 0.002 | 7 | -8 | 28 |
| | sulcus of corpus callosum | L | | 129.6 | 0.023 | 0.010 | 4.826 | 0.002 | -3 | -8 | 26 |
| | rostral cingulate zone | R | | 108.9 | 0.020 | 0.009 | 4.430 | 0.002 | 11 | -13 | 36 |
| | rostral cingulate zone, anterior to preSMA | L | | 200.4 | 0.021 | 0.010 | 4.340 | 0.002 | -5 | 16 | 33 |
| | superior frontal gyrus, medial surface, anterior | L, R | * R | 307.6 | 0.024 | 0.008 | 4.440 | 0.002 | -2 | 37 | 28 |
| | rostral cingulate zone, anterior/sulcus of corpus callosum | L | * | 110.6 | 0.020 | 0.008 | 4.736 | 0.002 | -8 | 22 | 20 |
| | dmPFC | R | | 79.5 | 0.023 | 0.012 | 4.283 | 0.003 | 5 | 53 | 29 |
| | vmPFC | L | | 162.4 | 0.024 | 0.013 | 4.431 | 0.003 | -2 | 32 | 5 |
| | frontal pole | R | | 155.5 | 0.019 | 0.006 | 4.424 | 0.002 | 12 | 67 | 12 |
| | caudate, head | R | | 81.2 | 0.022 | 0.011 | 4.555 | 0.002 | 11 | 14 | 4 |
| | cerebellum | R | * | 760.3 | 0.024 | 0.008 | 4.792 | 0.002 | 7 | -39 | -7 |
| | periacqueductal gray | | | 122.7 | 0.026 | 0.014 | 4.532 | 0.002 | -2 | -34 | -11 |
| | substantia nigra | L | | 146.9 | 0.024 | 0.011 | 4.297 | 0.003 | -4 | -13 | -9 |
| | medial geniculate nucleus | R | | 88.1 | 0.020 | 0.008 | 4.646 | 0.002 | 14 | -23 | -4 |
| **Actor identity** | mid temporal sulcus | L | | 102.0 | 0.027 | 0.014 | 4.588 | 0.002 | -49 | -43 | -10 |
| | SMG/parietal operculum | R | | 63.9 | -0.023 | 0.011 | -4.716 | 0.002 | 41 | -28 | 32 |
| | central sulcus/gyrus | R | | 70.8 | -0.024 | 0.015 | -4.299 | 0.003 | 20 | -21 | 48 |

| | | | | | | | | | | | |
|---|---|---|---|---|---|---|---|---|---|---|---|
| | posterior to PMv/White matter | R | * | 162.4 | -0.027 | 0.011 | -4.795 | 0.002 | 31 | -7 | 34 |
| **Non-emotion/Emotion** | central sulcus (M1) | L | | 81.2 | 0.027 | 0.013 | 4.367 | 0.003 | -31 | -20 | 49 |
| | middle frontal gyrus/superior frontal sulcus, adjacent to precentral sulcus, close to FEF/PMd | R | | 119.2 | 0.031 | 0.016 | 4.368 | 0.002 | 29 | 0 | 59 |
| | precuneus | L | | 79.5 | -0.028 | 0.016 | -4.367 | 0.002 | -14 | -53 | 55 |
| | superior frontal gyrus, medial surface | L, R | * | 95.0 | -0.028 | 0.015 | -4.848 | 0.002 | 1 | 24 | 52 |
| | caudal cingulate zone | L | | 63.9 | -0.023 | 0.014 | -4.383 | 0.002 | -10 | -5 | 45 |
| | thalamus | L | | 62.2 | -0.034 | 0.022 | -4.176 | 0.003 | -5 | -19 | 10 |
| **Implied Motion** | middle temporal sulcus | L | | 82.9 | -0.037 | 0.020 | -4.345 | 0.002 | -59 | -35 | -4 |
| | SPOC/IPSp | R | | 77.8 | 0.040 | 0.024 | 4.412 | 0.002 | 20 | -68 | 41 |
| | superior frontal sulcus, adjacent to FEF/PMd | L | | 67.4 | 0.034 | 0.020 | 4.384 | 0.002 | -22 | 1 | 58 |
| | PMv | L | | 152.1 | -0.035 | 0.017 | -4.360 | 0.002 | -43 | 7 | 34 |
| **Valence** | collateral sulcus/inferior parieto-occipital cortex | R | | 119.2 | -0.039 | 0.013 | -4.826 | 0.002 | 22 | -57 | 3 |
| | POC | R | * | 278.2 | -0.048 | 0.025 | -4.654 | 0.002 | 19 | -56 | 27 |
| | IPSa | R | * | 319.7 | -0.043 | 0.016 | -5.170 | 0.002 | 31 | -40 | 43 |
| | postcentral sulcus/SPL | R | | 140.0 | -0.042 | 0.019 | -4.193 | 0.003 | 13 | -36 | 58 |
| | SMG/parietal operculum | R | | 157.2 | -0.045 | 0.024 | -4.523 | 0.002 | 46 | -31 | 39 |
| | inferior frontal gyrus | R | * | 153.8 | -0.038 | 0.020 | -4.635 | 0.002 | 56 | 12 | 5 |
| | superior frontal sulcus, anterior. Next to the cluster of the motion RDM | R | | 93.3 | -0.039 | 0.013 | -4.827 | 0.002 | 19 | 31 | 43 |
| | lingual gyrus/cerebellum | R | | 98.5 | -0.048 | 0.031 | -4.308 | 0.002 | 7 | -68 | -5 |

**Table S4.** The univariate activation in the 10-action-category RDM clusters. The percent signal changes of each condition in each cluster were compared to the baseline (one-sample t test against 0).

| Reference RDM | Location | Hemisphere | survive cluster thresholding at initial p=.001 | Univariate ANOVA, p | Conditions showing activation to baseline |
|---|---|---|---|---|---|
| **Action categories** | cuneus | R | * | 0.481 | PH, TR, NE, AN, HA, SA |
| | hMT+/EBA | R | * | ***0.001** | all 10 conditions |
| | SPOC | R | | 0.065 | TR, AN, HA, SA |

| | | | | |
|---|---|---|---|---|
| IPSp | L | | 0.622 | all conditions except DW |
| IPSp | L | | 0.418 | DW, OD, TR, FE, AN, HA, SA |
| central sulcus (M1) | L | * | 0.376 | **None** |
| superior frontal sulcus | R | | 0.813 | AN |
| superior frontal sulcus, anterior cluster | R | | 0.299 | **None** |
| inferior frontal gyrus | R | | 0.776 | HA |
| precuneus | L | * | 0.866 | CH, DW, AN, HA |
| precuneus | R | | 0.327 | TR, NE, HA |
| precuneus | R | | 0.220 | DW, OD, TR, FE, AN, HA, SA |
| posterior cingulate gyrus | R | * | 0.132 | DW, FE |
| posterior cingulate sulcus | L | | 0.294 | CH, PH, TR, AN, HA |
| posterior cingulate sulcus/caudal cingulate zone | R | * | 0.377 | CH, OD, PH, TR, AN, HA |
| sulcus of corpus callosum | R | | 0.647 | CH, HA |
| sulcus of corpus callosum | L | | 0.434 | CH, OD, TR, NE, FE, AN, HA |
| rostral cingulate zone | R | | 0.752 | CH, DW, NE, FE, AN |
| rostral cingulate zone, anterior to preSMA | L | | 0.993 | all 10 conditions |
| superior frontal gyrus, medial surface, anterior | L, R | * R | 0.730 | **None** |
| rostral cingulate zone, anterior/sulcus of corpus callosum | L | * | 0.388 | DW, PH, AN |
| dmPFC | R | | 0.265 | PH |
| vmPFC | L | | 0.468 | **None** |
| frontal pole | R | | 0.649 | PH |
| caudate, head | R | | 0.569 | DW, TR, AN, HA |
| cerebellum | R | * | 0.550 | **None** |
| periacqueductal gray | | | 0.312 | OD, NE |
| substantia nigra | L | | 0.862 | PH, NE |
| medial geniculate nucleus | R | | 0.345 | OD, NE, AN, HA, SA |

**Table S5.** Clusters found in the searchlight RSA, for the perceived action and perceived emotion models (alpha=0.05, initial p=0.005, cluster size corrected). The R values were back transformed from Fisher's Z values. Subcortical and cerebellar clusters were listed after the cortical ones. * denotes clusters that survived thresholding at the initial p=.001. The average R values were in a range comparable to previous RSA studies (e.g. Fabbri et al., 2016; Tucciarelli et al., 2019).

| Reference RDM | Location | Hemisphere | initial p=.001 | cluster size (mm3) | average R | SD R | average t | average p | TAL x | TAL y | TAL z |
|---|---|---|---|---|---|---|---|---|---|---|---|
| **Perceived action** | mSTS | L | | 86.4 | 0.040 | 0.021 | 4.372 | 0.003 | -62 | -24 | -2 |
| | retrosplenial cortex | R | | 146.9 | -0.040 | 0.021 | -4.266 | 0.003 | 25 | -57 | 13 |
| | posterior collaterla sulcus, transversal | R | | 77.8 | -0.036 | 0.019 | -4.448 | 0.002 | 23 | -77 | -8 |
| | MGN | R | | 67.4 | -0.044 | 0.024 | -4.640 | 0.002 | 22 | -29 | 2 |
| | POC | R | | 138.2 | -0.045 | 0.021 | -4.538 | 0.002 | 16 | -58 | 28 |
| | superior frontal sulcus | R | | 72.6 | -0.056 | 0.034 | -4.604 | 0.002 | 16 | 7 | 46 |
| | frontal pole, medial superior frontal gyrus | R | | 72.6 | -0.047 | 0.024 | -4.375 | 0.002 | 16 | 50 | 14 |
| | SMA | R | | 77.8 | -0.046 | 0.027 | -4.277 | 0.003 | 8 | -21 | 59 |
| | dmPFC | L, R | | 197.0 | -0.050 | 0.023 | -4.395 | 0.002 | -1 | 42 | 43 |
| | CCZ | L | * | 437.2 | -0.055 | 0.026 | -4.482 | 0.002 | -5 | -9 | 44 |
| | RCZ | L | | 67.4 | -0.046 | 0.029 | -4.193 | 0.003 | -10 | 4 | 38 |
| **Perceived emotion** | PMd | L | | 119.2 | 0.041 | 0.019 | 4.671 | 0.002 | -34 | 3 | 52 |
| | SMG | R | | 76.0 | -0.035 | 0.017 | -4.575 | 0.002 | 59 | -29 | 34 |
| | SMG | R | | 82.9 | -0.034 | 0.015 | -4.458 | 0.002 | 52 | -34 | 29 |
| | SMG/aIPS | R | | 96.8 | -0.031 | 0.009 | -4.955 | 0.002 | 52 | -18 | -26 |
| | precentral gyrus, lateral | R | * | 133.1 | -0.032 | 0.012 | -4.820 | 0.002 | 49 | -5 | 45 |
| | claustrum | R | | 133.1 | -0.036 | 0.019 | -4.472 | 0.002 | 32 | -1 | 4 |
| | insula | R | | 65.7 | -0.036 | 0.014 | -4.508 | 0.002 | 29 | 22 | -5 |
| | putamen | R | * | 222.9 | -0.032 | 0.009 | -4.764 | 0.002 | 21 | 1 | -7 |
| | superior frontal gyrus | R | | 107.1 | -0.037 | 0.021 | -4.437 | 0.002 | 17 | 50 | 27 |
| | POC | R | | 191.8 | -0.031 | 0.011 | -4.601 | 0.002 | 14 | -58 | 35 |
| | cerebellum | R | | 98.5 | -0.035 | 0.017 | -4.147 | 0.003 | 14 | -54 | 13 |
| | superior frontal gyrus | R | | 69.1 | -0.037 | 0.022 | -4.102 | 0.003 | 14 | 57 | 5 |
| | precuneus | R | | 190.1 | -0.034 | 0.016 | -4.386 | 0.002 | 11 | -58 | 51 |
| | vmPFC | R | | 124.4 | -0.042 | 0.025 | -4.261 | 0.002 | 12 | 34 | -4 |
| | precuneus | R | | 67.4 | -0.040 | 0.022 | -4.398 | 0.002 | 12 | -44 | 56 |
| | dmPFC | R | * | 167.6 | -0.038 | 0.017 | -4.662 | 0.002 | 10 | 44 | 30 |
| | early visual areas | R | * | 159.0 | -0.036 | 0.012 | -4.647 | 0.002 | 7 | -77 | -2 |
| | medial superior frontal gyrus | R | | 96.8 | -0.033 | 0.016 | -4.366 | 0.002 | 3 | 25 | 50 |
| | callosal sulcus | R | | 373.2 | -0.037 | 0.015 | -4.444 | 0.002 | 1 | -19 | 28 |
| | dmPFC | L | | 74.3 | -0.034 | 0.016 | -4.391 | 0.002 | -7 | 52 | 28 |
| | cuneus | L | | 67.4 | -0.037 | 0.021 | -4.535 | 0.002 | -16 | -72 | 17 |
